# iCliff Taylor’s version: Robust and Efficient Activity Cliff Determination

**DOI:** 10.1101/2025.03.09.642269

**Authors:** Kenneth López-Pérez, Ramón Alain Miranda-Quintana

**Affiliations:** Department of Chemistry and Quantum Theory Project, University of Florida, Gainesville, FL 32611, USA

**Keywords:** similarity, Tanimoto, iSIM, activity cliff

## Abstract

Activity cliffs represent an important challenge to tackle in cheminformatics and drug design. One of the most common indicators to quantify them is the SALI index. Here we expose mathematical limitations of SALI’s formulation, the most evident: it is undefined in instances where the similarity between two molecules is one. We show how using a simple Taylor’s series can aid this main problem, yielding a defined expression that can capture the ranking information from the original SALI. The second issue to solve is the quadratic complexity of using SALI to describe the roughness of the activity landscape of a set. Here, we propose iCliff, an indicator that can quantify the roughness in linear complexity. For this, we leverage the iSIM framework to obtain the average similarity of the set and a rearrangement to obtain the average of the squared property differences. The calculations for 30 different AC-focused databases suggest that there is a strong correlation between iCliff and the average pairwise of SALI’s pairwise Taylor Series. To further explore the individual effects of removing each molecule in the activity landscape, we propose complementary iCliff. With this tool, we were able to identify the molecules that have a high number of activity cliffs with the rest of the molecules in the set.

## 1. INTRODUCTION

One problem that medicinal chemists and cheminformaticians have pointed out for decades are Activity Cliffs (ACs).^1^ ACs are defined as compounds with high similarity but with a large difference in bioactivity.^1–4^ ACs break the similarity principle: *“similar molecules have similar properties”*^5^; in Figure 1 we include an example of a pair of molecules that do not follow it. Since much of the reasoning behind drug design often relies on the similarity principle, Acs are of special attention in Quantitative Structure Activity-Relationship (QSAR) studies, making them a current challenge to overcome.^2,6^With the growth of the use of Machine Learning (ML) models for QSAR models, there is still high a necessity to identify/quantify activity cliffs efficiently, and how to mitigate their influence on a model’s performance.^7,8^ The first step towards identifying ACs is defining what are similar compounds. A plethora of similarity approaches directly applied to ACs have been proposed. One of the alternatives is Matched Molecular Pairs (MMPs)^9^ which defines a similar pair of compounds as two molecules that have a structural modification only on one site. Another way is simply getting pairs from an analog series.^10–12^ Arguably the classical and most common way of quantifying similarity is the use of a similarity index like the popular Tanimoto index.^13,14^ These indexes are calculated from binary fingerprints or molecular descriptors. One of the advantages of Tanimoto, or any other similarity index, is that we get a numerical value that tells us how similar the molecules are. However, there is no stoned-set agreement on what threshold value should be used to call a pair of molecules similar since the values are highly dependent on the representation.^15^ The second step is to assess the activity differences, in the field a 100-fold difference has been commonly used^2,3^, but this number does not take into consideration that potency distributions might vary depending on the target.^16^

**Figure 1.**
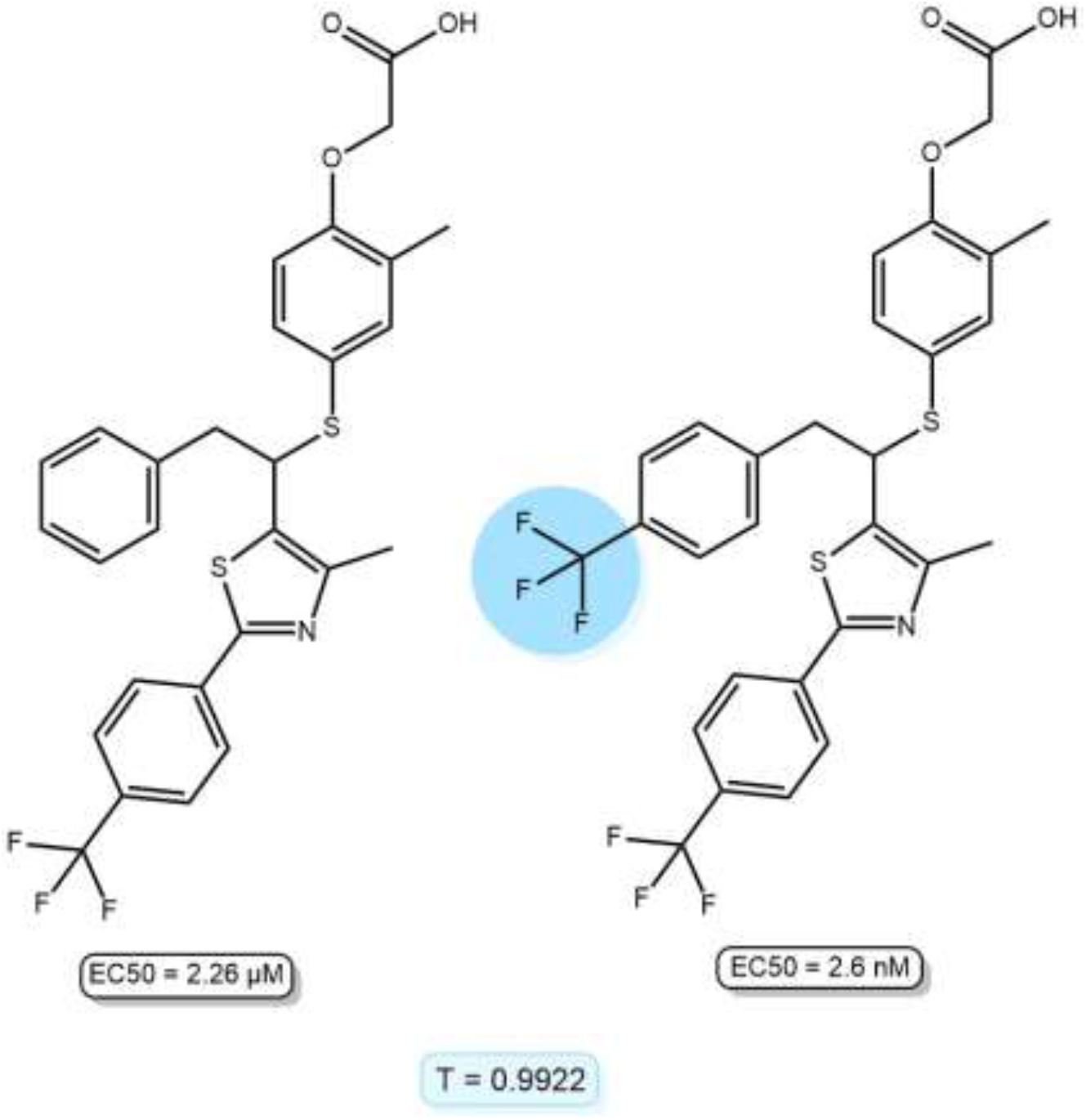
Activity cliff example for a pair of molecules with peroxisome proliferator-activated receptor delta (PPARδ) activity.^8^ Tanimoto similarity calculated from RDKIT binary fingerprints (2048 bits).

Ways of integrating similarity and activity differences for ACs identification include Structure-Activity Similarity (SAR) maps, where the similarities are plotted against the potency differences, yielding an easily interpretable plot to identify ACs.^17^ The Structure-Activity Landscape Index (SALI) calculates the magnitude of the property change with respect to the distance (1 – similarity) of two compounds.^18^ The SALI is defined in Equation 1, there, *P*_*x*_ stands for the value of property *P* for molecule *x*, and *s*_*ij*_ is the similarity between two molecules *i* and *j* (while there are multiple ways to quantify this similarity, it is often just calculated using the Tanimoto index).

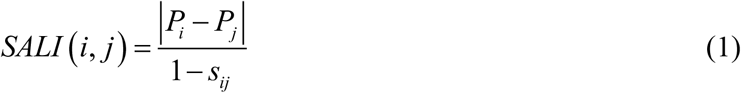

Unlike the pairwise nature of SALI, other approaches have focused on the development of metrics that quantify the overall roughness of the activity landscapes. For example, the Roughness Index (ROGI)^19^ is a metric based on the change of property dispersions in clusters after performing hierarchical clustering on the distance matrix with a range of thresholds. One of the limitations of ROGI is the O(*kN*^*2*^) complexity due to the need for the pairwise distance matrix.^19^ The Structure-Activity Relationship Index (SARI) is an approach that employs a continuity score (property-weighted pairwise similarity) and a discontinuity score (average potency differences between pairs with higher Tanimoto than 0.6 multiplied by the Tanimoto similarity).^20^ SARI standardizes the scores on 16 reference datasets, relying on user-defined parameters/data. Also, it scales O(*N*^*2*^).^20^ The eSALI index is a global roughness index that uses the extended similarity^21,22^ framework to get a global similarity value for the set and the differences between properties and the average.^23^ Despite the linear complexity of this method, O(*N*), eSALI requires a user-defined similarity threshold to calculate the extended similarity.^23^ Large-scale Machine Learning models have been reported for the prediction of activity cliffs classification of pairs (AC or non-AC pairs); previous identification with other methods is required to train the models.^24^

Here, we present an innovative way to quantify the presence of ACs, free of some of the key issues of traditional approaches like the necessity for user-defined parameters, mathematical undefinition, and complex scalability. Moreover, we show with the use of iSIM techniques we can calculate a global activity-landscape roughness metric and also identify molecules with a prevalence of ACs, which we exemplify over a varied set of libraries.

## 2. THEORY

While very popular due to its simplicity and ease of calculation, the SALI indicator has three key weaknesses:

1. It is unbounded.
2. It is undefined in instances where *s*_*ij*_ = 1.
3. It requires considering every possible pair of molecules in a set, so identifying potential molecules with ACs demands O(*N*^2^) computational effort.

In this section, we will propose ways to modify Eq. (1) such as to overcome these issues, while keeping its advantages.

First, we note that even if the *s*_*ij*_ = 1 condition could be relatively rare, it cannot be guaranteed to never occur, especially if the molecules are represented using binary fingerprints. In those cases, it could be possible that different compounds are encoded in the same way, depending on the resolution power of the chosen binary encoding. However, even beyond these clashes, very similar molecules will produce denominators that are close to zero, making Eq. (1) difficult to interpret and also prone to numerical instabilities. Since the root of this issue is the formulation of the traditional SALI as a fraction with a denominator that could be arbitrarily close (if not identical) to zero, the recipe to solve this problem is quite simple: we must reformulate Eq. (1) as a product instead of as a division. The key to doing this is the following Taylor expansion:

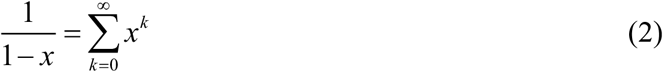

So, the Taylor Series (TS) for SALI can be written as:

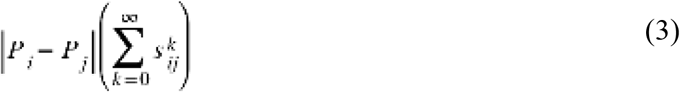

The second problem associated with the SALI, namely, the O(*N*^2^) demand to calculate it for all the pairs of molecules in a set, could be approached by using (*P*_*i*_ − *P*_*j*_)^2^ instead of |*P*_*i*_ − *P*_*j*_| to capture property differences, and the recently proposed iSIM (instant similarity)^25^ formalism to calculate the average similarity of the molecules of the set.

For pairs of molecules, the Taylor Series SALI (TS_SALI) can be truncated into any desired finite term for it to be useful. After changing the absolute value difference for the squared difference, we explore three possibilities, truncating at *k* = 1, 2, and 3, respectively. In these cases, and purely for the sake of having normalized values for the TS_SALI, we include normalization constants as:

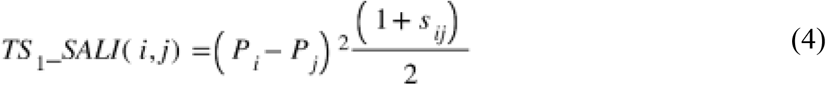

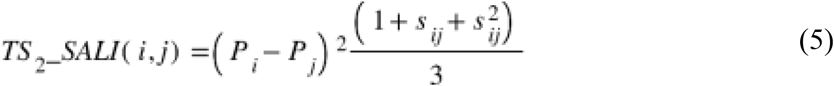

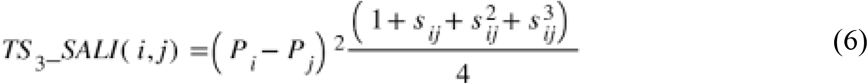

Now to obtain a set-wise indicator we can calculate the average of squared differences between properties in a set by decomposing it into individual terms as the following:

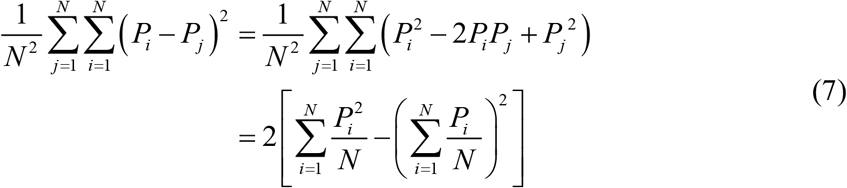

and we can easily see from the r.h.s. that this can be calculated in O(*N*) time.

iSIM greatly facilitates handling the similarity component of this expression, since it directly gives access to the average of all the pairwise similarities for any given library. For the particular case of the Tanimoto index, the iSIM Tanimoto (iT) for *N* molecules encoded in *M*-bits fingerprints is just:

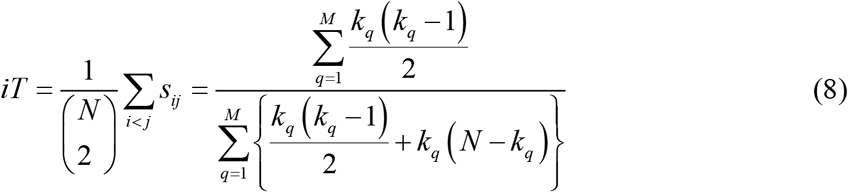

where *k*_*q*_ is the sum of all the elements of the *q*^th^ column of the matrix of fingerprints.

In the traditional sense, the roughness of the activity-property landscape can be directly linked to the average of the SALI or TS_SALI over all pairs of compounds in the set. With the exposed ingredients, we can now get an expression that will measure the roughness of the activity landscape for a set of *N* molecules in O(*N*) time. Inspired by the previously reported eSALI, we present iCliff in Equation 9 with k =3 (other truncations are possible). Higher iCliff values mean a higher presence of ACs in a set.

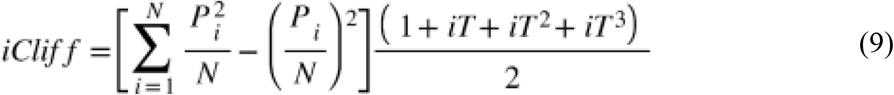

Now that we have an indicator for the whole set, we want to obtain a metric that can tell us if a molecule is likely to have ACs with the rest of the molecules in the set. In short, we need to evaluate the impact of removing a molecule from the set, both on the property differences and the overall similarity of the set. For this, we recur to the concept of complementary similarity. That is, the iSIM of the set when the *i*^th^ molecule is removed from the set, 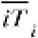. This process has also O(*N*) complexity since for the iSIM calculation we only need the linear sum of fingerprints from a set, we can easily subtract the *i*th fingerprint and recalculate iSIM with *N-1* molecules.^25^ The complementary similarity value will tell us how different or similar from the rest of the set is that molecule.

For the properties this is quite simple to do: if we pre-calculate 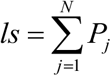 and 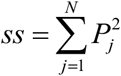, then:

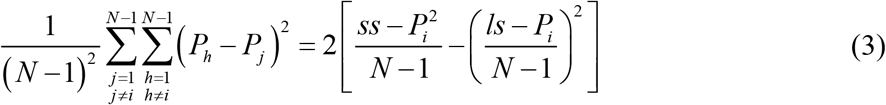

With these ingredients, we can define the complementary iCliff, 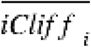, for the *i*^th^ molecule as a way to encapsulate the impact of removing a single molecule on the full activity-property landscape:

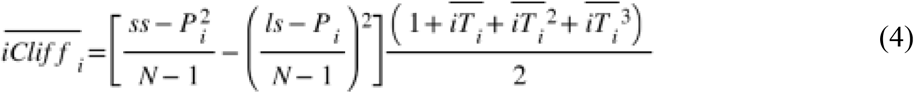

These modifications are critical to finding an efficient way to identify molecules that could potentially present ACs. Low complementary iCliff values indicate that the molecule roughens the activity landscape of the set.

## 3. COMPUTATIONAL METHODS AND SYSTEMS

### Data

We used CHEMBL datasets for 30 different molecular targets, specially curated to focus on the study of Activity Cliffs. SMILES and EC50/Ki were sourced from van Tilborg et al.^8^ (dataset sizes and codes included in SI, Table S1) Molecules were represented with RDKIT^26^ (2048 bits), MACCS^27^ (166 bits), and ECFP4^28^ (1024 bits) binary fingerprints, computed with RDKIT.^26^

Properties, *K*_i_ or EC50, (originally reported in nM) were converted into M, and applied negative logarithm to calculate the pK_i_ or pEC50 were calculated. Properties were normalized using Min-Max normalization, for a set of properties *P* = {*P*_1_, *P*_2_, …, *P*_3_} the normalization is as follows:

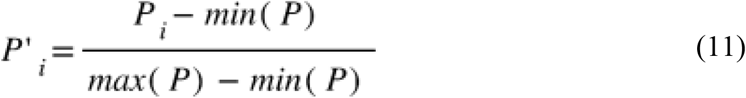

### Methods

For all the databases and fingerprints, pairwise matrices of SALI and TS_SALI (1^st^, 2^nd^, and 3^rd^ truncations) were calculated as defined in Equations 1, 4, 5, and 6; respectively. Additionally, the pairwise SALI matrix with squared property difference instead of absolute value was also computed. Ranking correlation between matrices was measured with Kendall’s Tau coefficient. ^29^

iCliff values for all databases with ECFP4 fingerprints were calculated following equation 9, before and after removing pairs of ACs identified from the TS_SALI matrices. Complementary iCliff was calculated for each molecule in the databases, and correlation analysis between them and the column-wise sum of the TS_SALI matrices was done with Kendall’s tau and Jaccard^30^ of fractions of the sets.

All scripts used in this work are available at https://github.com/mqcomplab/iCliff.

## 4. RESULTS

The first thing that we checked is the agreement between SALI and TS_SALI. Here, we do not pay too much attention to the absolute SALI and TS_SALI values, but to the relative rankings that they induce in the pairs of molecules. To quantify this we calculated the full SALI(i, j) and TS_SALI(i, j) matrices (and, for the sake of consistency and due to the above-mentioned SALI issues, we defined the SALI(i, i) = 0 and did not considered in the analysis pairs with undefinitions in the SALI), “flattened” them and calculated Kendall’s tau (Kτ) value between the resulting one-dimensional vectors. As shown in Fig. 2A), for all the libraries and fingerprints considered, we see the expected increase in Kτ going from 1^st^ to 2^nd^ to 3^rd^ order truncation of the Taylor series in the TS_SALI expressions. Given that there is virtually no extra computational cost in moving from the 1^st^ to the 3^rd^ order, we recommend using the 3^rd^ order truncation as the *de facto* standard to calculate the TS_SALI.

**Figure 2.**
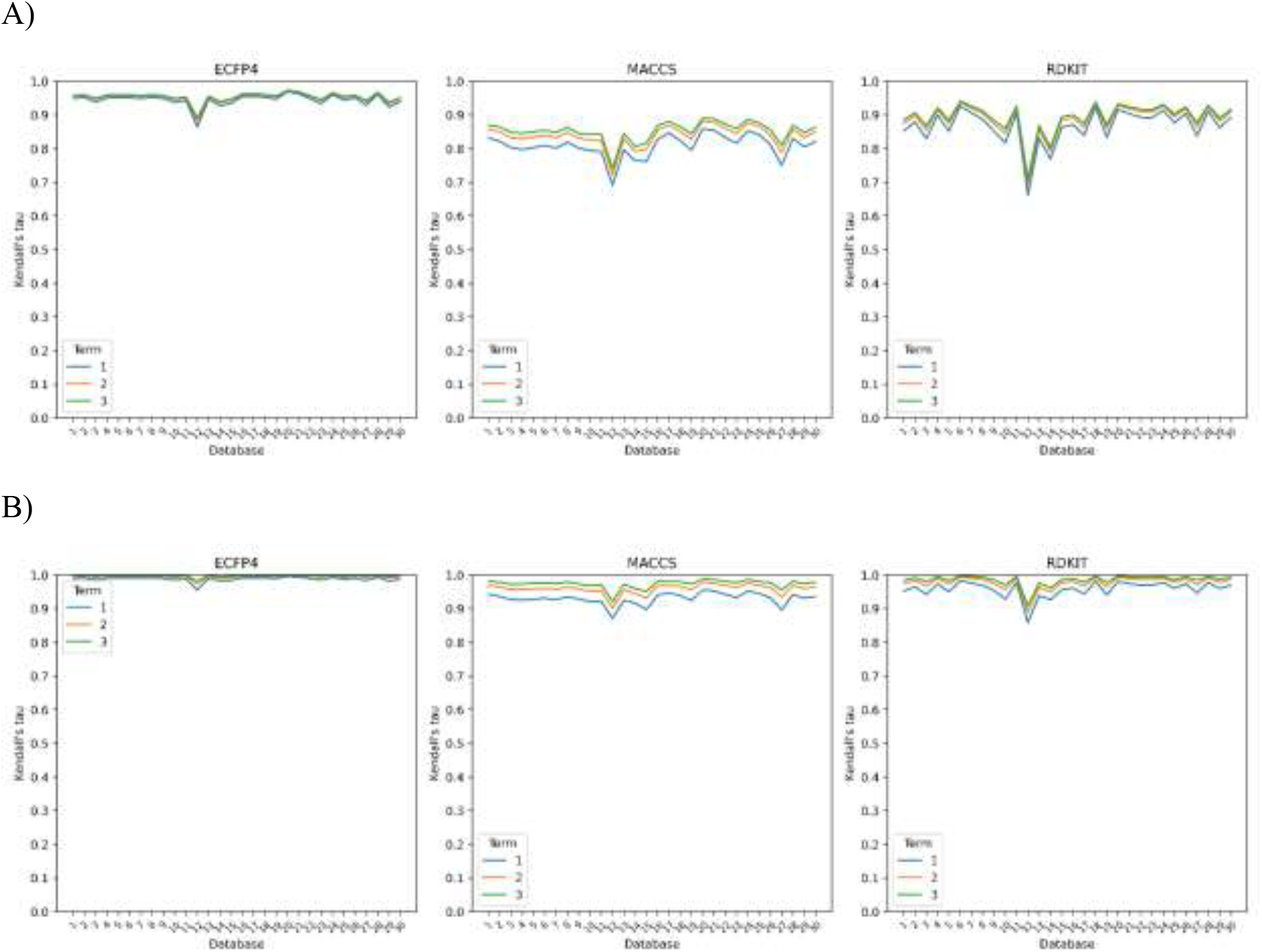
Kendall’s tau correlations between the SALI and TS_SALI values for different truncation orders over the 30 studied libraries represented with ECFP4 (1024 bits), MACCS (166 bits), and RDKIT (2048 bits) binary fingerprints. In A) SALI is defined as in Eq. 1, in B) we use SALI with the squared property difference.

Another interesting trend is found in the analysis of how the correlations change with the fingerprint type. Note that, consistently, the ECFP representation gives better results than the RDKit fingerprints, which in turn surpasses the MACCS keys. Remarkably, there are cases where 1^st^ order ECFP is even better than 3^rd^ order RDKit and MACCS. Even more, in most cases, ECFP at 3^rd^ order shows Kτ values > 0.95. We also studied the correlation between the TS_SALI matrices and a modified version SALI using the squared property difference instead of the absolute value. In Fig 2B), we can see how the ranking correlations are even better, suggesting that most of the discrepancies between SALI and TS_SALI are not attributed to the Taylor Series expansion. Either way, the Kendall τ values for ECFP4 fingerprints are the most consistent with SALI, so we will continue our analysis focusing on those fingerprints.

Having established an agreement between SALI and TS_SALI, we now move to an analysis of the TS_SALI values. In Figure 3 we can see the distributions of TS_SALI values for two databases, the rest of the histograms are included in the Supporting Information. The shape of the distribution is expected, previous reports mention that the prevalence of activity cliffs is usually low, which agrees with the skewness towards zero of the obtained distributions. Lower TS_SALI values mean less of an activity cliff the pair of molecules is. Overall, from the distributions we can deduct a threshold value to define an AC pair. The TS_SALI value for the 95th percentile is 0.11 (on average for the 30 databases) and 0.16 for the 99th percentile. Since in the literature, the prevalence of ACs is between 1-5%^2,4^, we suggest that a good cutoff to define activity cliffs with TS_SALI would be around 0.15. Again, no threshold is set in stone, it will vary with representation, user necessities, and target. The ChEMBL_2835 database (12^th^ database in the plots, the one that correlates the worst between SALI and TS_SALI values, and the smallest one) has an atypical distribution compared to the other databases, it is noticeable that the number of activity cliffs (TS_SALI > 0.15) is greater than the distributions for the rest of the databases, all similar to the ChEMBL_264 distribution shown in Fig 3A).

**Figure 3.**
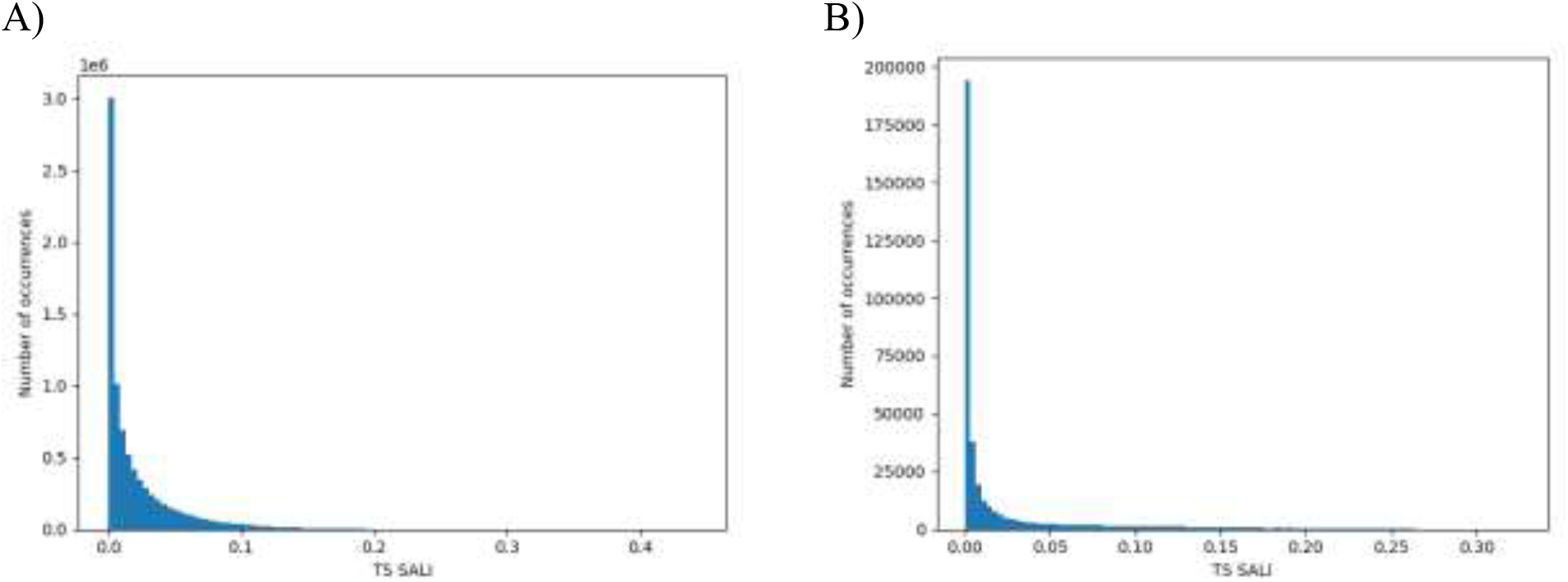
Distribution of pairwise TS_SALI values using 3^rd^ order truncation for the A) ChEMBL264 and B) ChEMBL2835 libraries represented with ECFP4 fingerprints.

Now we move to the iCliff approach, a method to quantify the activity landscape roughness/smoothness without the need to compute the pairwise TS_SALI matrix. First, from the global level point of view, we calculate the iCliff for each database and compare it with the average TS_SALI matrix for each database. In Figure 4, we can see that for almost all the databases there is a numerical agreement between iCliff and the mean value of TS_SALI. Despite the clear deviation of one database, the correlation and ranking correlation are strong and positive, with both the determination coefficient and Kendall’s τ above 0.90. The most deviated point corresponds to the database with the lowest number of molecules and high prevalence of ACs (according to Fig. 3B distribution).

**Figure 4.**
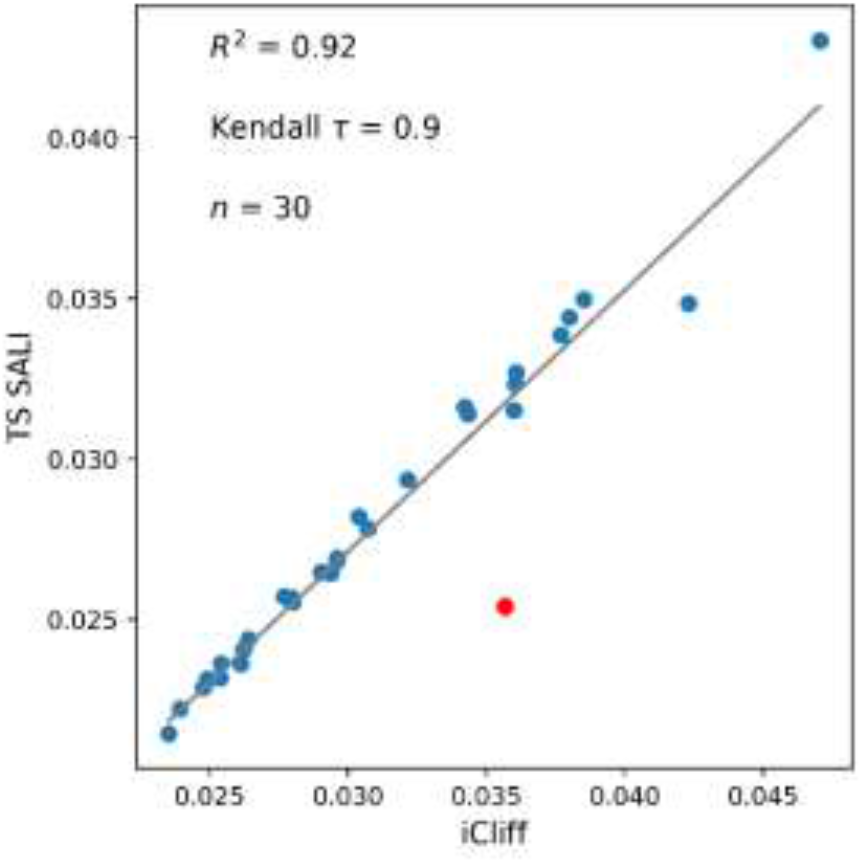
Relationship between the average TS SALI value (from the TS SALI pairwise matrix) and the iCliff value for the 30 ChEMBL’s databases represented with ECPF4 fingerprints (1024 bits). The tendency line is shown in gray.

The higher the iCliff value the rougher the landscape of the set is. To prove that in fact our method is good at describing the overall roughness of the activity landscape, we calculate the iCliff for the complete databases, and then the iCliff after removing the molecules in pairs with TS_SALI higher than the 99^th^ percentile. Depending on the database the number of removed molecules does not directly translate to a fixed percentage of the data since in some cases molecules could have ACs with multiple other molecules. In Figure 5, we can see how there is a decrease in the iCliff value after removing the molecules in ACs, proving that the proposed method accomplishes the goal of describing the activity landscape of the sets.

**Figure 5.**
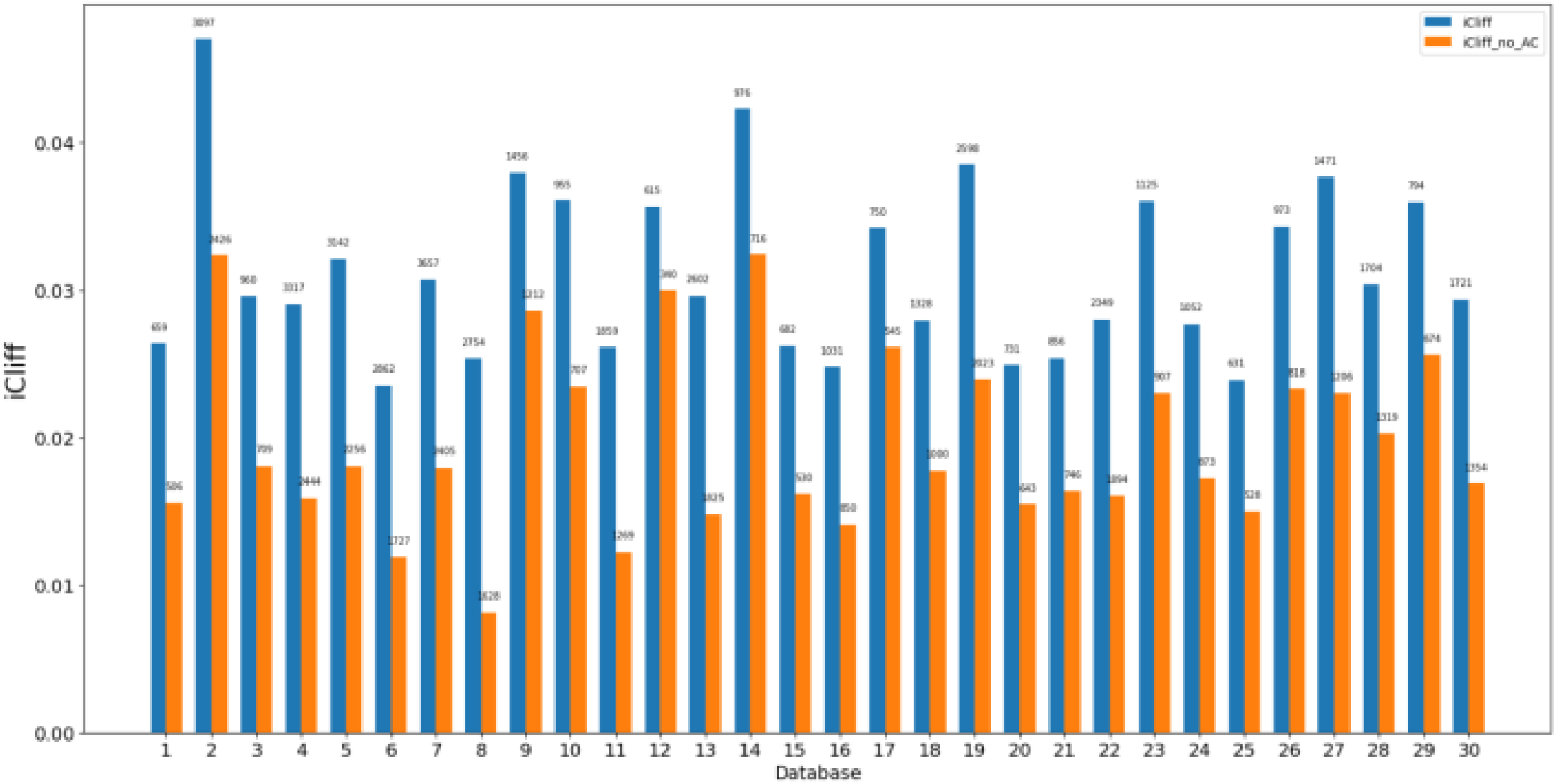
iCliff values for the ChEMBL databases studied before and after removing activity cliffs (molecules in pairs with TS_3__SALI > the 99^th^ percentile). Molecules represented ECFP4 (1024-bit) fingerprints. The number of molecules annotated over the bars.

Having established the numerical and ranking correlation between iCliff and the average of the TS SALI matrix and showcasing its purpose; we proceed to identify individual molecules that are present in AC pairs with complementary iCliff. Our strategy is to correlate the O(*N*) complementary iCliff results, Eq. (4), with the column-wise summation of the TS SALI O(*N*^2^) matrix (*cTS_SALI*). Molecules that will be present in one relevant or several activity cliffs, will have high values for the column-wise sum, as it will include the information of all the computations with the rest of the molecules. To explore this connection, we first analyzed the Kτ between these indicators. Given the results discussed previously in this section, here we only focus on the versions of these indices that use a 3^rd^ order Taylor truncation, and ECFP fingerprints. Note in Fig. 6 that the overall results show a great consistency between the *cTS_SALI* matrix sum and iCliff. In 29 out of 30 cases, the Kτ is above 0.8, only one value is below 0.8 (the CHEMBL2835 library, the same library that did not correlate well in Fig. 3).

**Figure 6.**
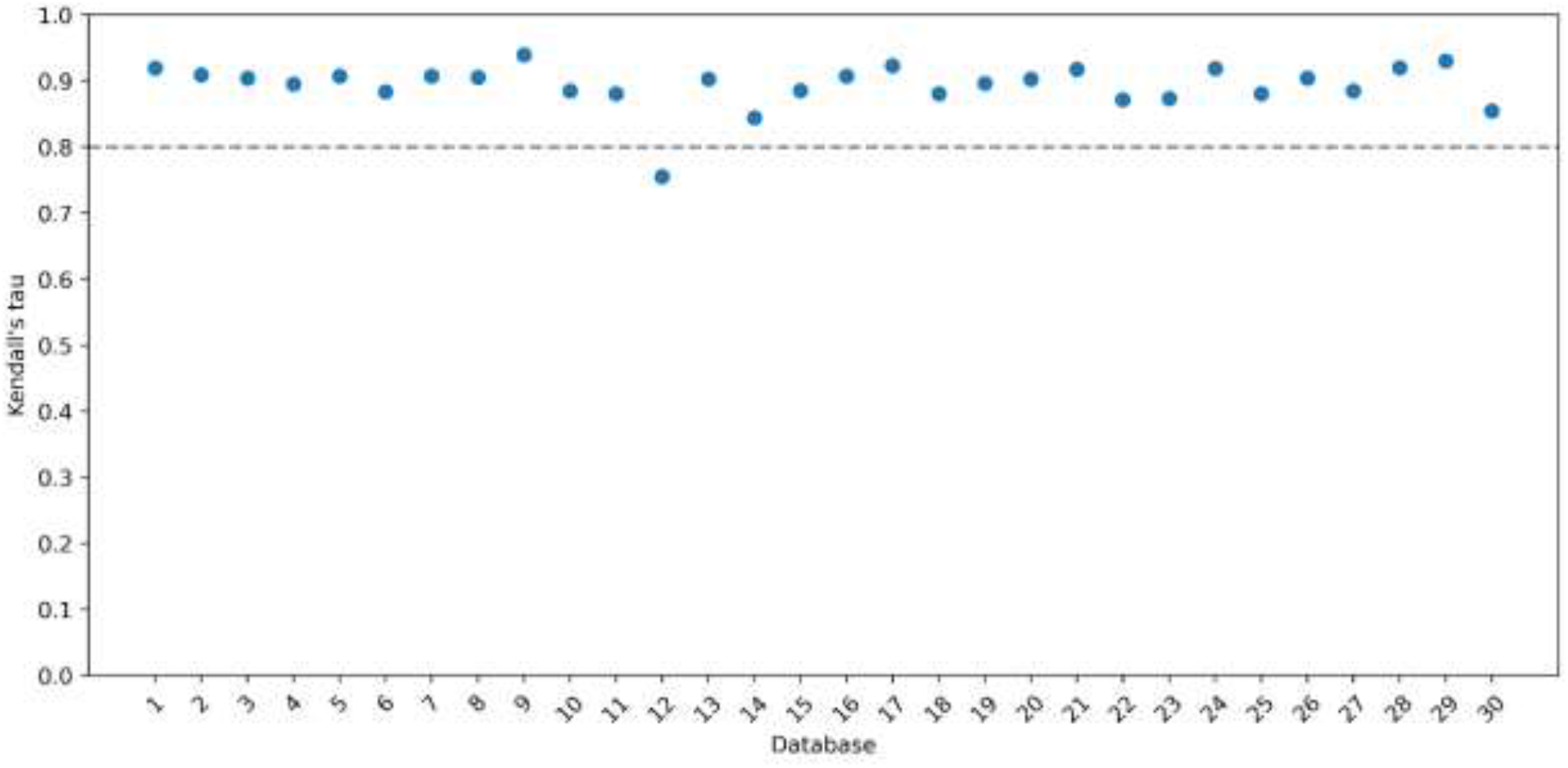
Kendall’s tau correlations between the cTS_SALI and iCliff values over the 30 studied libraries using ECFP4 fingerprints.

However, this test is more demanding than what is actually needed in practice. The Kτ between the column-wise TS SALI and iCliff takes into account the order of all possible molecules in the set, even if those molecules are not actively participating in ACs. While it is reassuring that even in those very strict conditions these two indicators agree to such an extent, in practice, one could be more interested in merely just identifying the top compounds with the biggest influence in the structure-activity landscape. That is, it is important to see how well iCliff can identify exactly the same molecules as *cTS_SALI* when we only want to explore a fraction of the library. To study this, we use the (set) Jaccard similarity, to check the degree of coincidence between the top *k* molecules find by *cTS_SALI* and complementary iCliff, *J*_*k*_. In short, we look at:

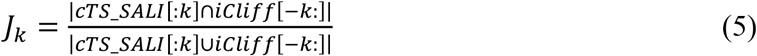

Here, |X| is the size of set X, and the “Pythonic” [:*k*] [-*k*:] notation indicates that we are selecting the first or last *k* elements (respectively) of the given ordered lists. (The difference in first or last elements for the ordered *cTS_SALI* and complementary iCliff lists stands from the previously mentioned inverse ordering of these indicators). Notice in Fig. 7 that even the most “problematic” set (CHEMBL2835_Ki) shows excellent performance in identifying the top 10% (vertical line in Fig. 7) molecules roughening the structure-activity landscape. The ranking does not match after this point for this database, but this supports the ability of complementary iCliff to identify the molecules that have activity cliffs with others and also explains the deviations in the previous analyses. For example, the CHEMBL1862_Ki library shown in Fig. 7 almost immediately settles at *J*_*k*_ > 0.8, with many instances even above 0.9, thus indicating an excellent agreement between complementary iCliff and *cTS_SALI*. This tendency can be observed in all the other libraries (SI), where the Jaccard values quickly stabilize after a short period of instability in picking the first handful of molecules.

**Figure 7.**
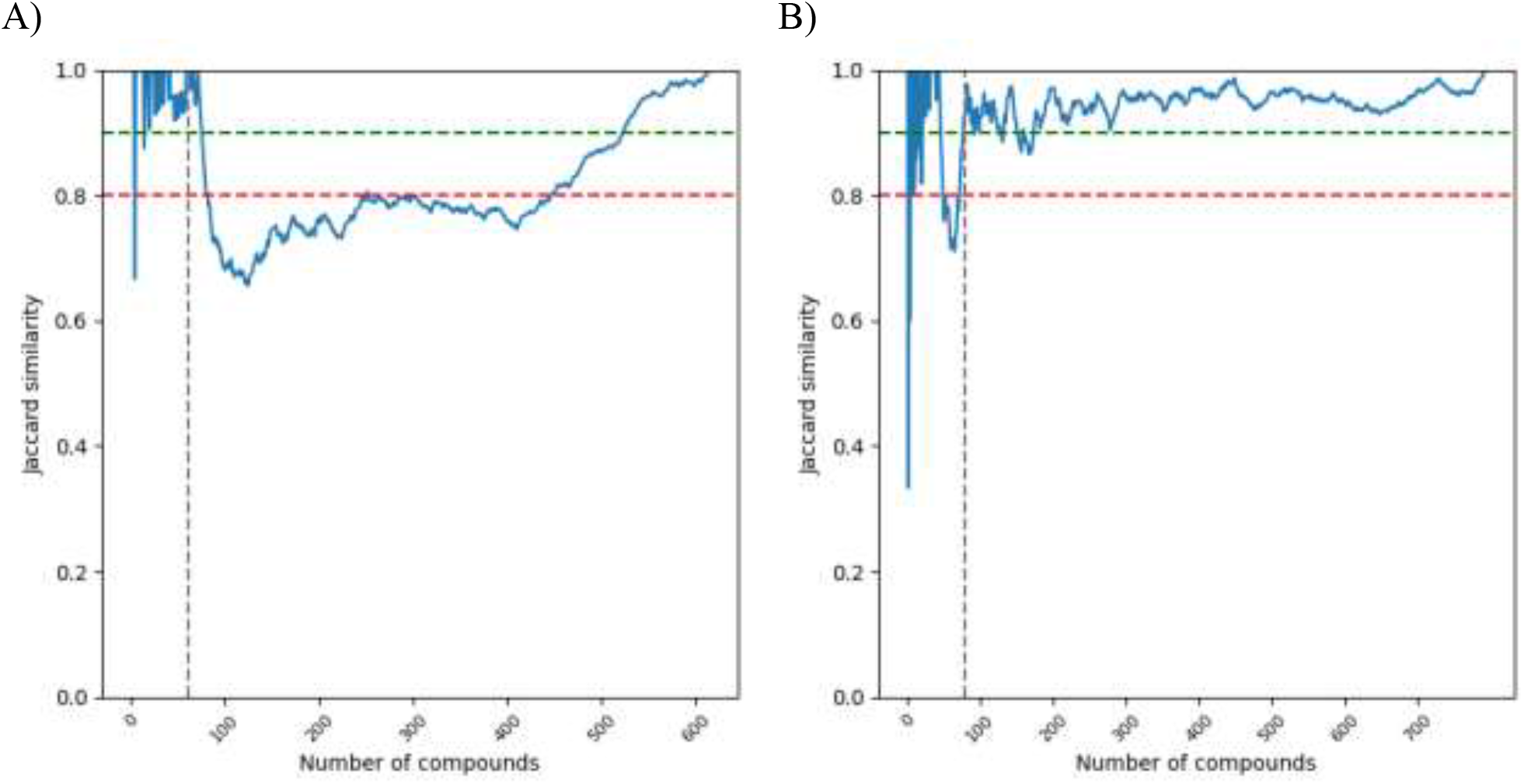
Variation of the Jaccard similarity between the ranking of cTS_SALI and complementary iCliff, when varying the fraction of molecules. A) CHEMBL2835_Ki and B) CHEMBL1862_Ki. Horizontal lines at 0.8 and 0.9, vertical line at 10% of the size of each set.

Finally, to look at how removing the 5% identified molecules with complementary iCliff affects the structure-activity landscape, we calculated the iCliff before and after removing them (similarly to the shown results in Figure 5). Here in Figure 8, we can see how in all cases the activity landscape is softened after removing the molecules identified with low complementary iCliff. Hence, iCliff can be a tool for tasks where a flat landscape is needed, i.e. ML models. We also want to point out the dramatic drop in the iCliff value for the 12^th^ database, this means that the “problematic” molecules, had activity cliff with high number molecules, when removed the iCliff value gets really close to zero.

**Figure 8.**
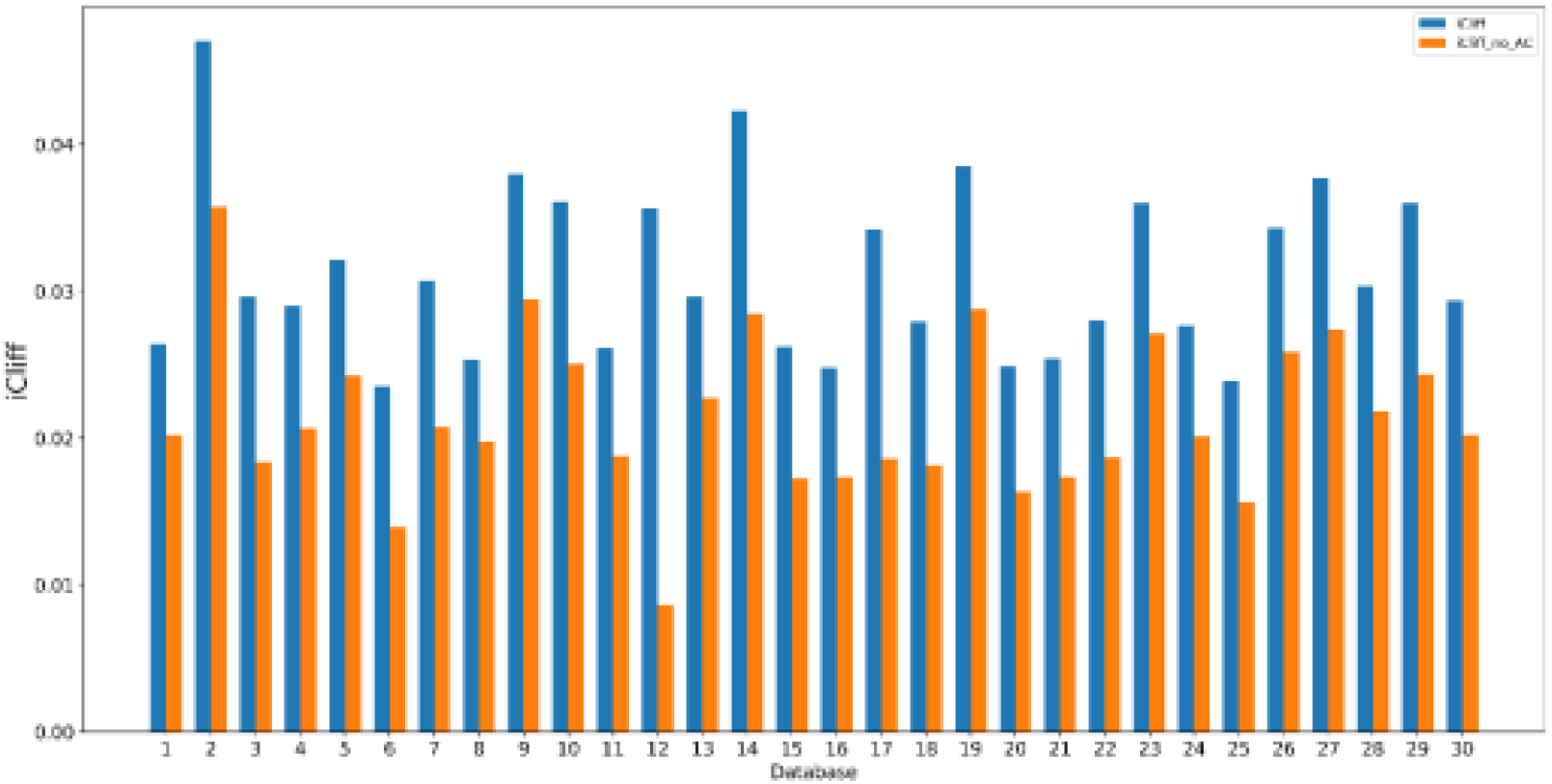
iCliff values for the studied ChEMBL databases before and after removing activity cliffs (5% of molecules with lowest complementary iCliff). Molecules are represented with ECFP4 (1024 bit) fingerprints.

## 5. CONCLUSIONS

We have presented both a new, robust, way to quantify the structure-activity landscape of a set and a faster method to identify molecules in a dataset that could present this behavior. First, we introduce TS_SALI, we show how one can bypass the notorious zero-division errors in the traditional SALI formulation. The simple substitution of a low-order truncation of a Taylor series is enough to solve this issue while keeping most of the ranking information contained in SALI. Even a 1^st^ order approximation gives excellent performance across the board for all the studied libraries and fingerprint types. However, given the inexpensive nature of the low-order corrections, and the considerable improvement in the Kt values, we recommend using the 3^rd^ order truncation. Also, in all the considered cases, the TS_SALI values have a better ranking correlation with SALI when using ECFP fingerprints, consistently better than RDKIT and MACCS counterparts.

Additionally, we introduced iCliff, a global metric to quantify the structure-activity landscape roughness. In the analysis of 30 AC-focused databases, we show how there is a strong positive correlation between the proposed iCliff and the average of the pairwise TS_SALI matrix. Finally, we showed how to identify the molecules with the biggest impact on the topography of the structure-activity landscape, with the introduction of the complementary iCliff indicator. Complementary iCliff combines the complementary similarity notion of iSIM (describing the impact that a molecule has in the overall similarity of the set) with a more computationally friendly way to quantify the impact of removing a property value from the pool of molecules, as a way to gauge the individual contribution of a compound to the ACs in the library. Remarkably, complementary iCliff provides this information demanding just O(*N*) time and memory, without the need to exhaustively explore all possible pairs of molecules, as in traditional SALI-based methods. The general agreement between iCliff and the column-wise sum of the TS_SALI matrix is excellent in almost all the studied cases, but it is even more impressive when tasked with identifying the top fraction of molecules with the biggest role in shaping the structure-activity landscape. Overall, we have shown that is possible to not only globally characterize these landscapes, but also to zoom in on their local properties much more efficiently than previously thought. We anticipate that these techniques will open the door to other efficient ways to study structure-activity relations.

## ACKNOWLEDGEMENTS

We thank support from the National Institute of General Medical Sciences and the National Institutes of Health under award number R35GM150620.

## DATA AVAILABILITY STATEMENT

The code used to calculate iCliff can be found here: https://github.com/mqcomplab/iCliff.

