## Supplementary Information for "iCliff Taylor’s version: Robust and Efficient Activity Cliff Determination"

**Table 1**. Studied databases names, codes, and sizes. Sourced from van Tilborg et al.[1]

| Code | Name | n |
|---|---|---|
| 1 | CHEMBL1871_Ki | 659 |
| 2 | CHEMBL244_Ki | 3097 |
| 3 | CHEMBL4005_Ki | 960 |
| 4 | CHEMBL214_Ki | 3317 |
| 5 | CHEMBL233_Ki | 3142 |
| 6 | CHEMBL264_Ki | 2862 |
| 7 | CHEMBL234_Ki | 3657 |
| 8 | CHEMBL204_Ki | 2754 |
| 9 | CHEMBL2147_Ki | 1456 |
| 10 | CHEMBL237_EC50 | 955 |
| 11 | CHEMBL219_Ki | 1859 |
| 12 | CHEMBL2835_Ki | 615 |
| 13 | CHEMBL237_Ki | 2602 |
| 14 | CHEMBL2971_Ki | 976 |
| 15 | CHEMBL4616_EC50 | 682 |
| 16 | CHEMBL218_EC50 | 1031 |
| 17 | CHEMBL2034_Ki | 750 |
| 18 | CHEMBL287_Ki | 1328 |
| 19 | CHEMBL236_Ki | 2598 |
| 20 | CHEMBL4203_Ki | 731 |
| 21 | CHEMBL262_Ki | 856 |
| 22 | CHEMBL235_EC50 | 2349 |
| 23 | CHEMBL3979_EC50 | 1125 |
| 24 | CHEMBL238_Ki | 1052 |
| 25 | CHEMBL2047_EC50 | 631 |
| 26 | CHEMBL231_Ki | 973 |
| 27 | CHEMBL4792_Ki | 1471 |
| 28 | CHEMBL228_Ki | 1704 |
| 29 | CHEMBL1862_Ki | 794 |
| 30 | CHEMBL239_EC50 | 1721 |

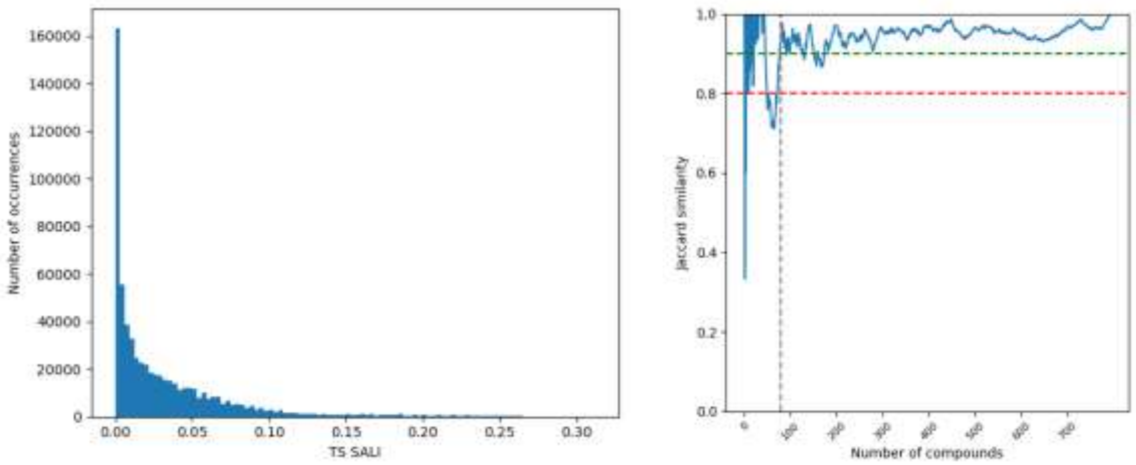


**Figure 1.** Distribution of pairwise TS_SALI values using 3rd order truncation. (Left) Variation of the Jaccard similarity between the ranking of cTS_SALI and complementary iCliff for database CHEMBL1862_Ki. (right).

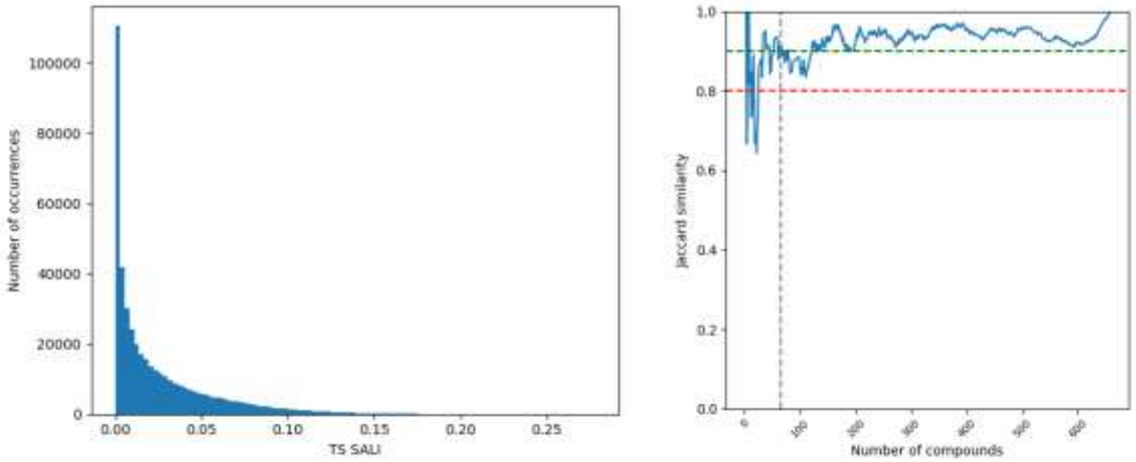


**Figure 2.** Distribution of pairwise TS_SALI values using 3rd order truncation. (Left) Variation of the Jaccard similarity between the ranking of cTS_SALI and complementary iCliff for database CHEMBL1871_Ki. (right).

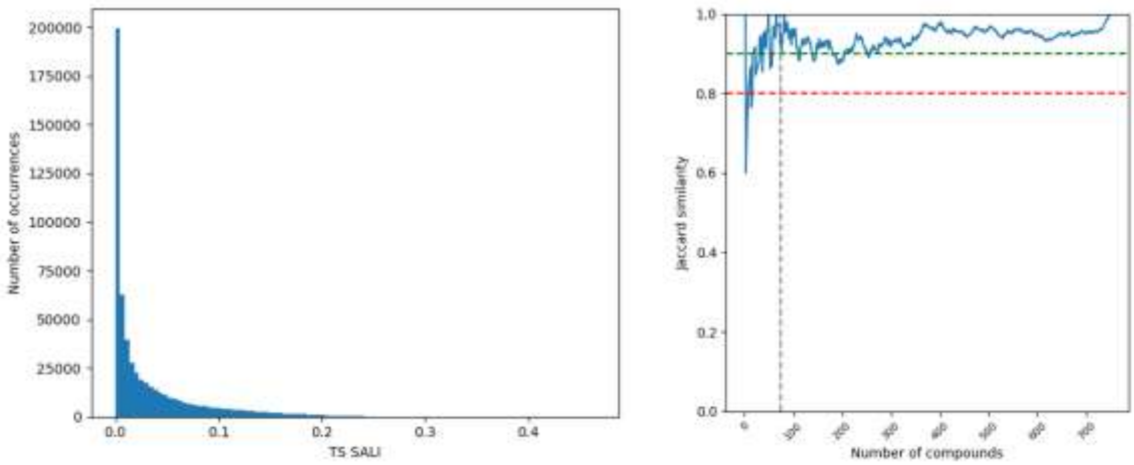


**Figure 3.** Distribution of pairwise TS_SALI values using 3rd order truncation. (Left) Variation of the Jaccard similarity between the ranking of cTS_SALI and complementary iCliff for database CHEMBL2034_Ki. (right).

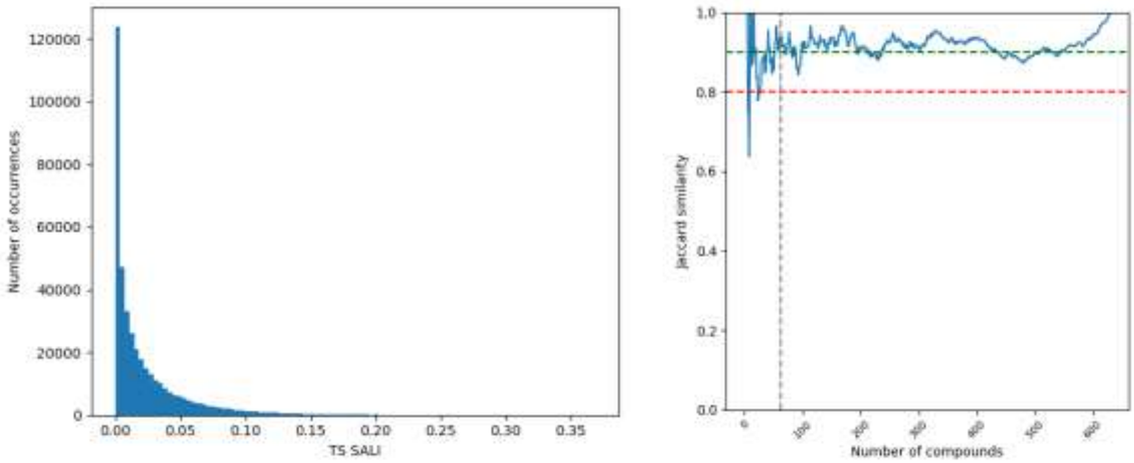


**Figure 4.** Distribution of pairwise TS_SALI values using 3rd order truncation. (Left) Variation of the Jaccard similarity between the ranking of cTS_SALI and complementary iCliff for database CHEMBL2047_EC50. (right).

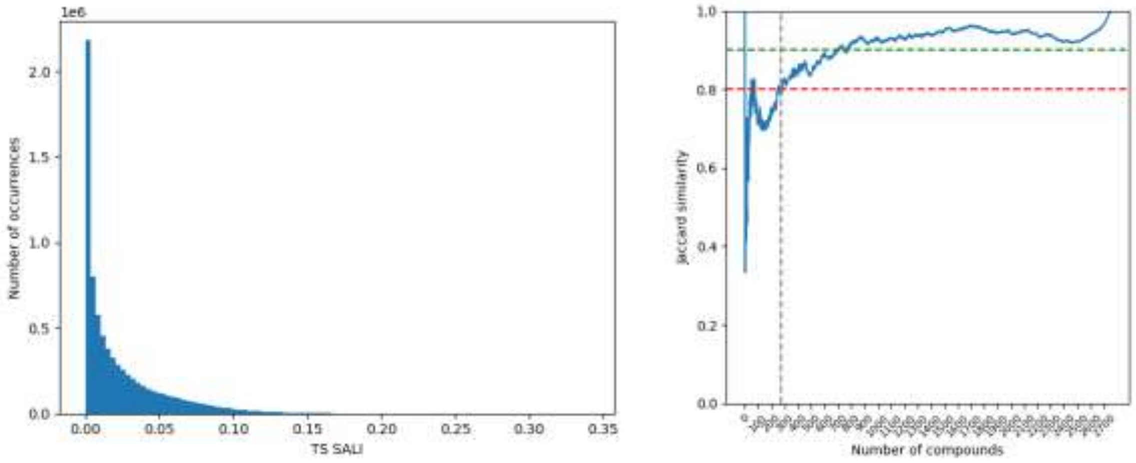


**Figure 5.** Distribution of pairwise TS_SALI values using 3rd order truncation. (Left) Variation of the Jaccard similarity between the ranking of cTS_SALI and complementary iCliff for database CHEMBL204_Ki. (right).

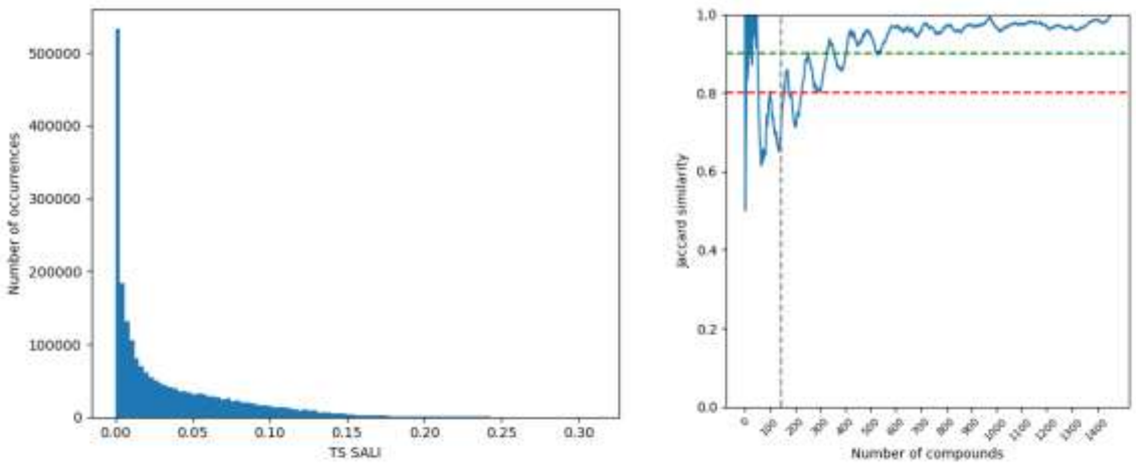


**Figure 6.** Distribution of pairwise TS_SALI values using 3rd order truncation. (Left) Variation of the Jaccard similarity between the ranking of cTS_SALI and complementary iCliff for database CHEMBL2147_Ki. (right).

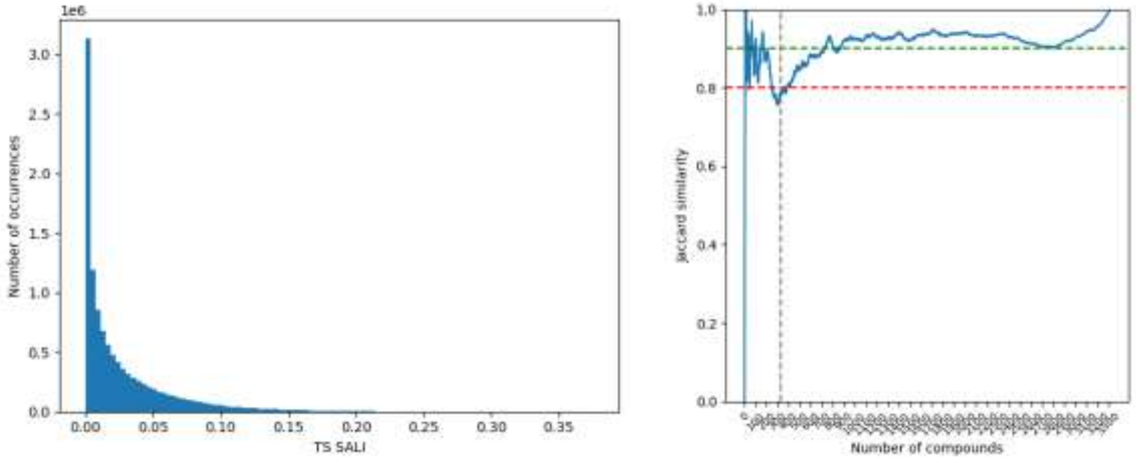


**Figure 7.** Distribution of pairwise TS_SALI values using 3rd order truncation. (Left) Variation of the Jaccard similarity between the ranking of cTS_SALI and complementary iCliff for database CHEMBL214_Ki. (right).

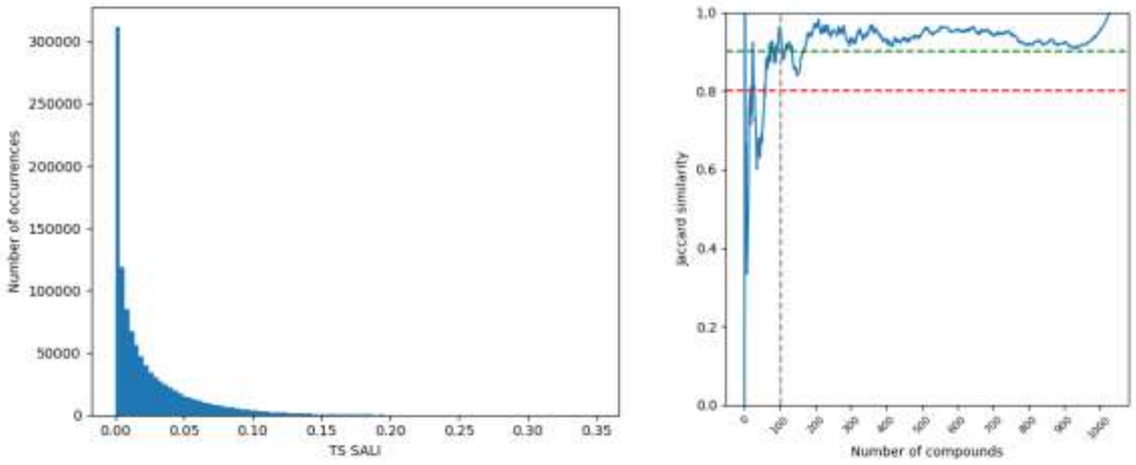


**Figure 8.** Distribution of pairwise TS_SALI values using 3rd order truncation. (Left) Variation of the Jaccard similarity between the ranking of cTS_SALI and complementary iCliff for database CHEMBL218_EC50. (right).

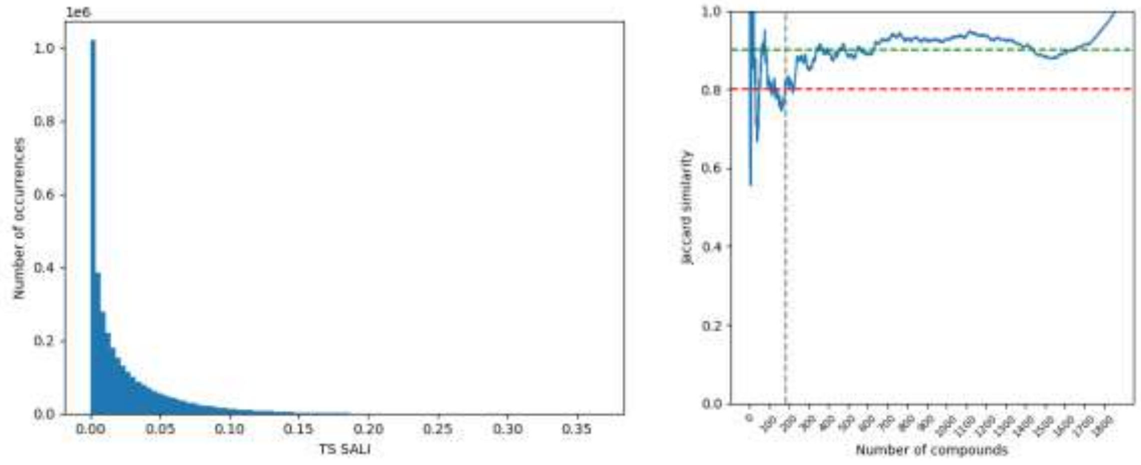


**Figure 9.** Distribution of pairwise TS_SALI values using 3rd order truncation. (Left) Variation of the Jaccard similarity between the ranking of cTS_SALI and complementary iCliff for database CHEMBL219_Ki. (right).

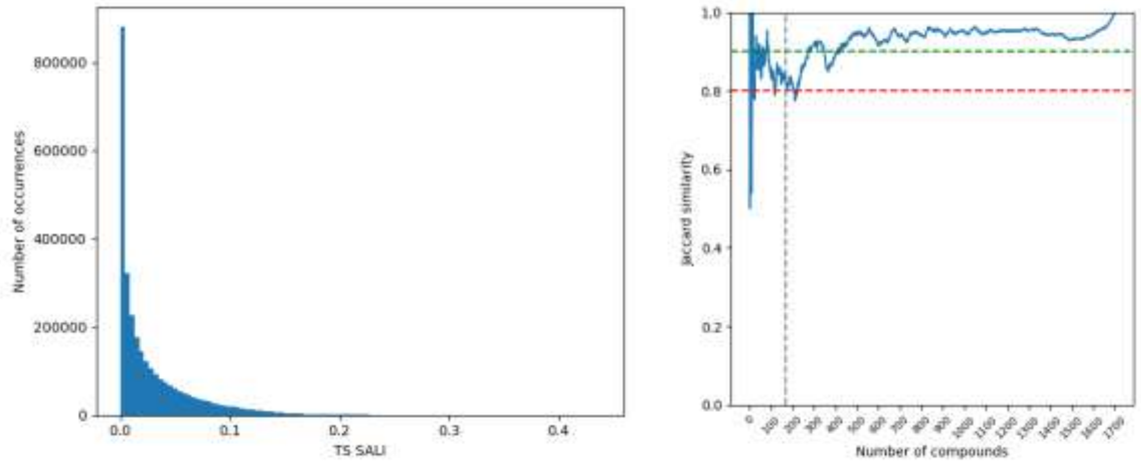


**Figure 10.** Distribution of pairwise TS_SALI values using 3rd order truncation. (Left) Variation of the Jaccard similarity between the ranking of cTS_SALI and complementary iCliff for database CHEMBL228_Ki. (right).

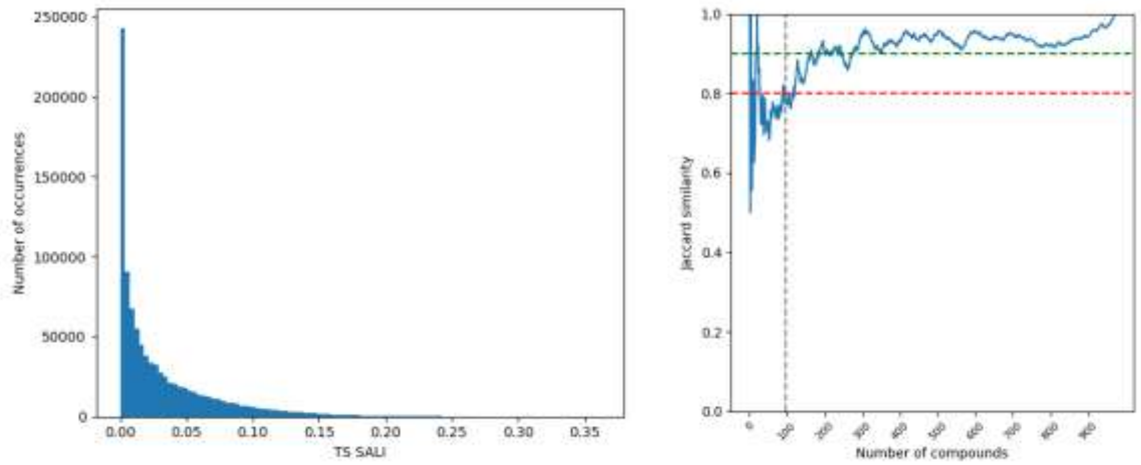


**Figure 11.** Distribution of pairwise TS_SALI values using 3rd order truncation. (Left) Variation of the Jaccard similarity between the ranking of cTS_SALI and complementary iCliff for database CHEMBL231_Ki. (right).

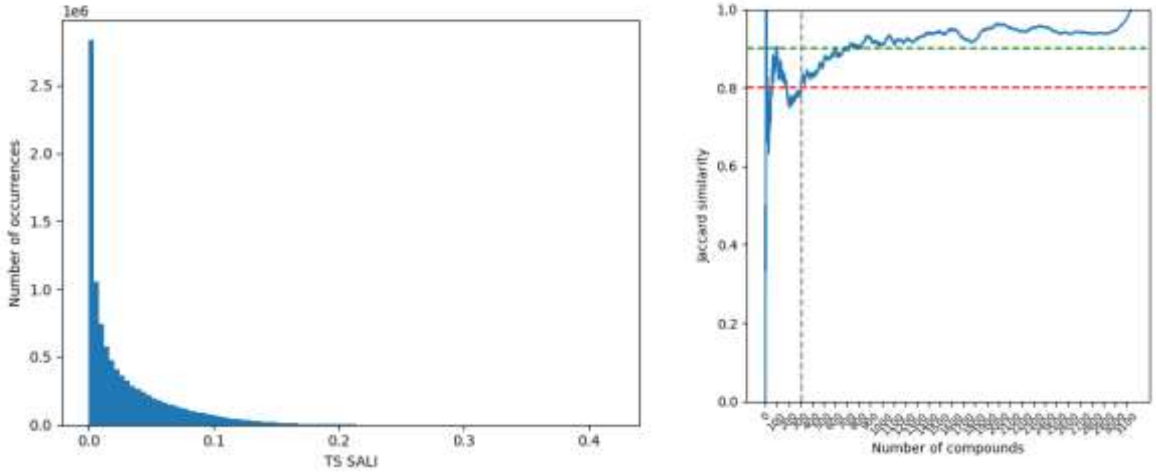


**Figure 12.** Distribution of pairwise TS_SALI values using 3rd order truncation. (Left) Variation of the Jaccard similarity between the ranking of cTS_SALI and complementary iCliff for database CHEMBL233_Ki. (right).

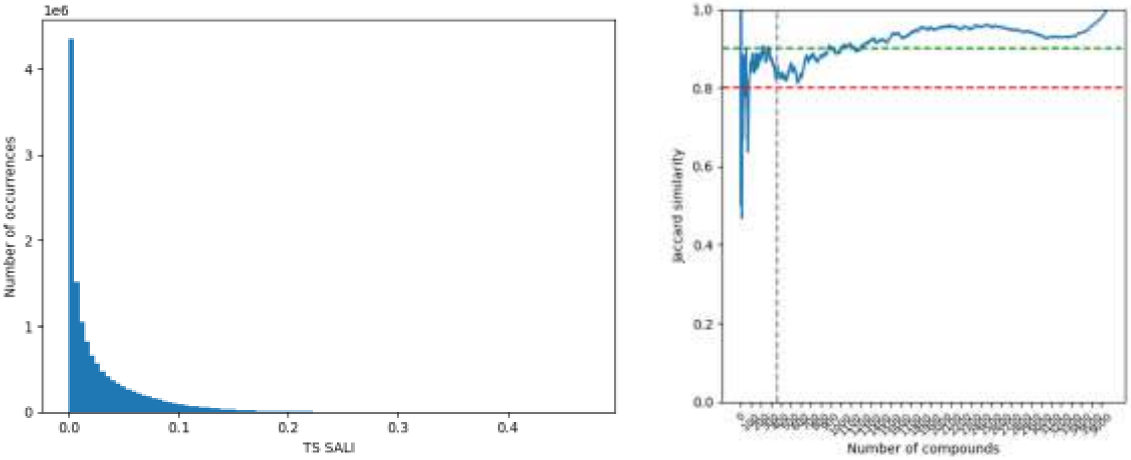


**Figure 13.** Distribution of pairwise TS_SALI values using 3rd order truncation. (Left) Variation of the Jaccard similarity between the ranking of cTS_SALI and complementary iCliff for database CHEMBL234_Ki. (right).

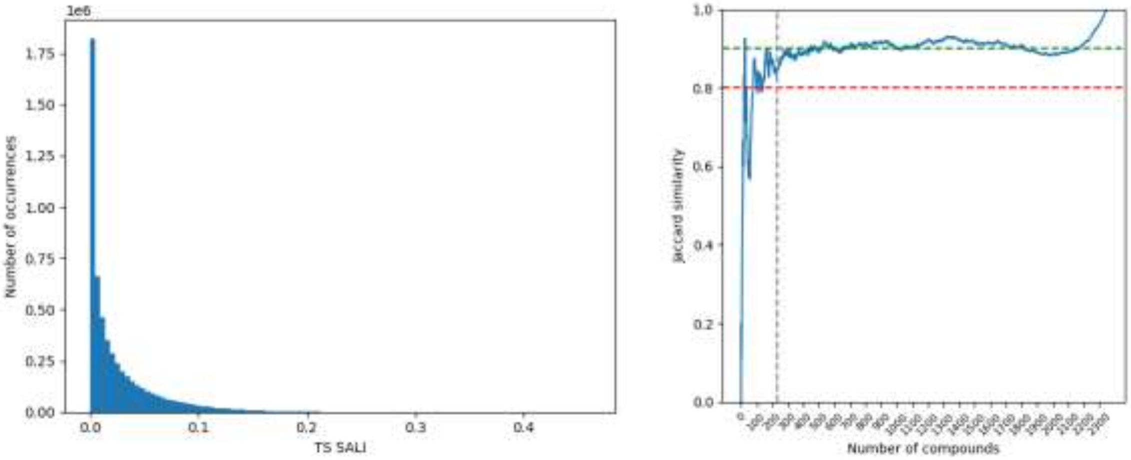


**Figure 14.** Distribution of pairwise TS_SALI values using 3rd order truncation. (Left) Variation of the Jaccard similarity between the ranking of cTS_SALI and complementary iCliff for database CHEMBL235_EC50. (right).

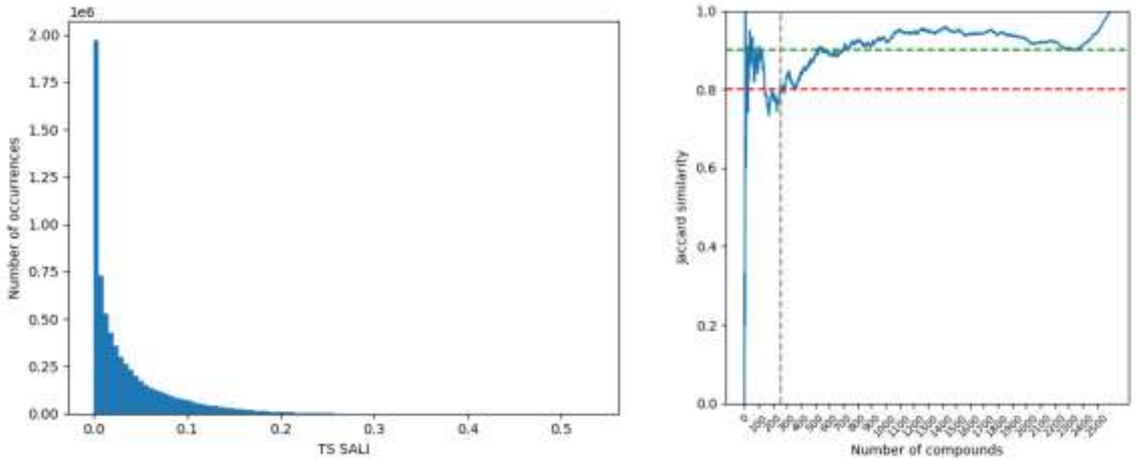


**Figure 15.** Distribution of pairwise TS_SALI values using 3rd order truncation. (Left) Variation of the Jaccard similarity between the ranking of cTS_SALI and complementary iCliff for database CHEMBL236_Ki. (right).

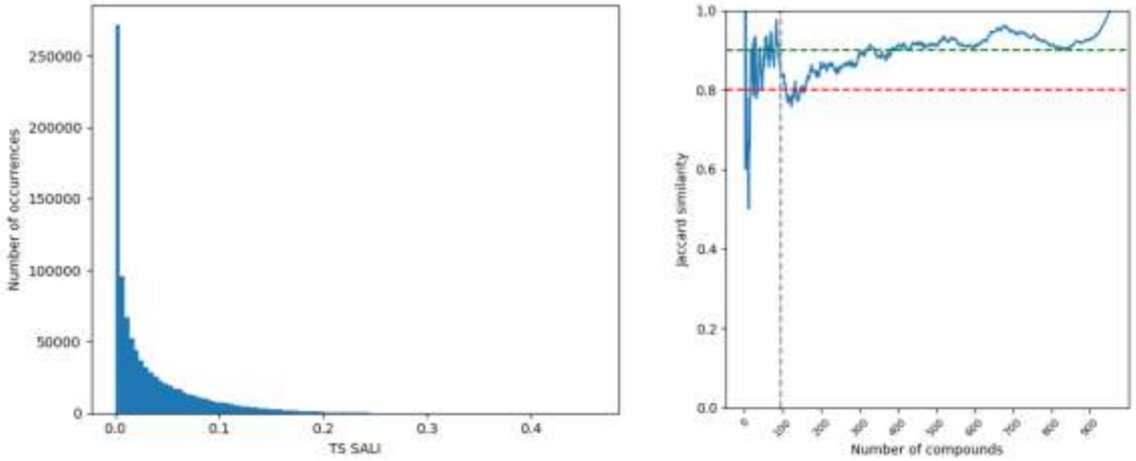


**Figure 16.** Distribution of pairwise TS_SALI values using 3rd order truncation. (Left) Variation of the Jaccard similarity between the ranking of cTS_SALI and complementary iCliff for database CHEMBL237_EC50. (right).

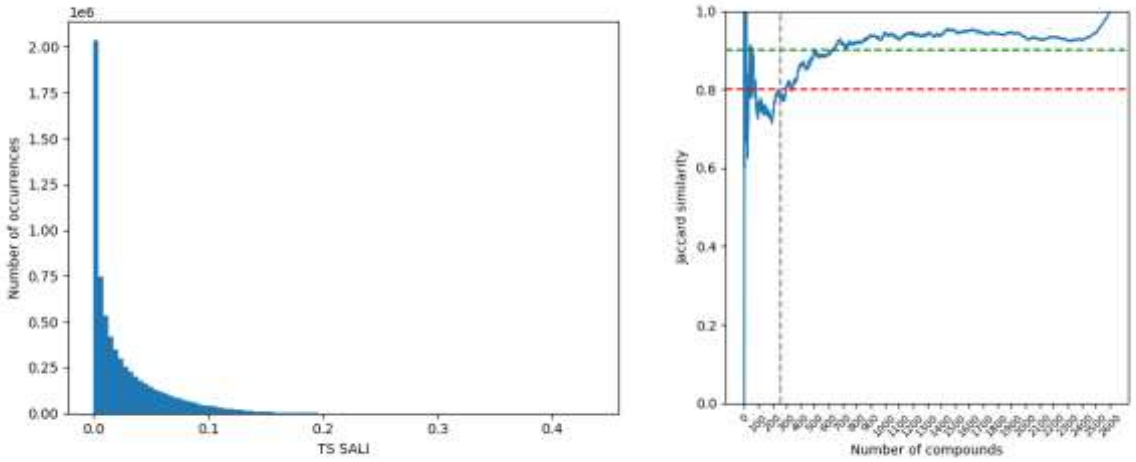


**Figure 17.** Distribution of pairwise TS_SALI values using 3rd order truncation. (Left) Variation of the Jaccard similarity between the ranking of cTS_SALI and complementary iCliff for database CHEMBL237_Ki. (right).

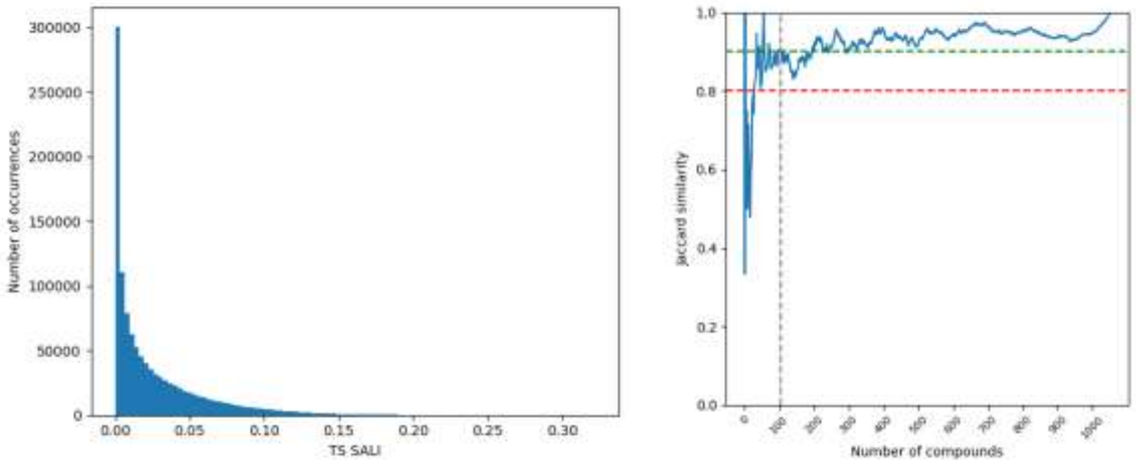


**Figure 18.** Distribution of pairwise TS_SALI values using 3rd order truncation. (Left) Variation of the Jaccard similarity between the ranking of cTS_SALI and complementary iCliff for database CHEMBL238_Ki. (right).

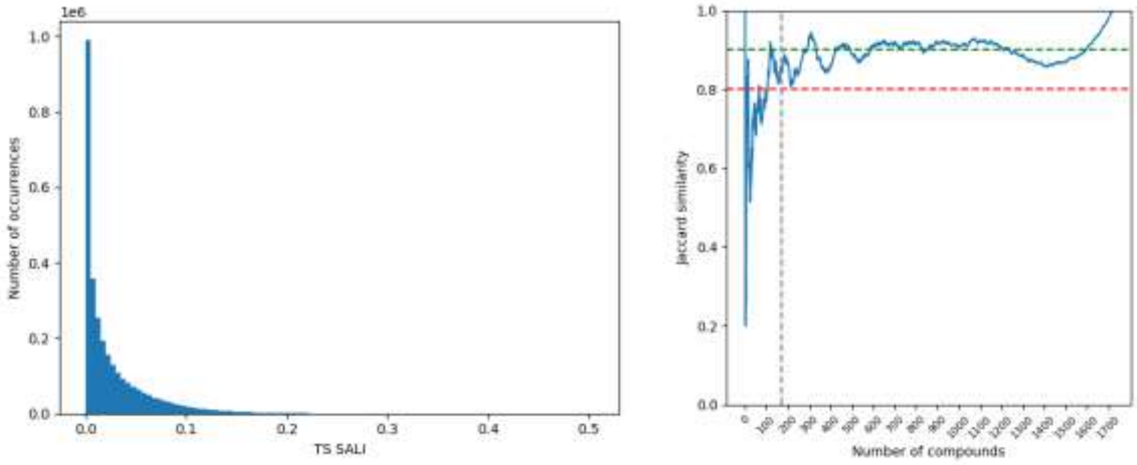


**Figure 19.** Distribution of pairwise TS_SALI values using 3rd order truncation. (Left) Variation of the Jaccard similarity between the ranking of cTS_SALI and complementary iCliff for database CHEMBL239_EC50. (right).

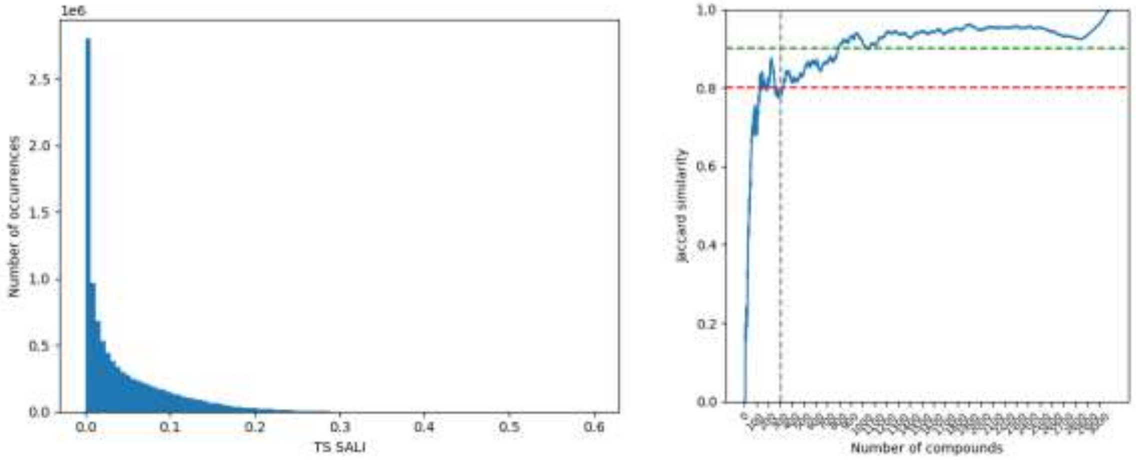


**Figure 20.** Distribution of pairwise TS_SALI values using 3rd order truncation. (Left) Variation of the Jaccard similarity between the ranking of cTS_SALI and complementary iCliff for database CHEMBL244_Ki. (right).

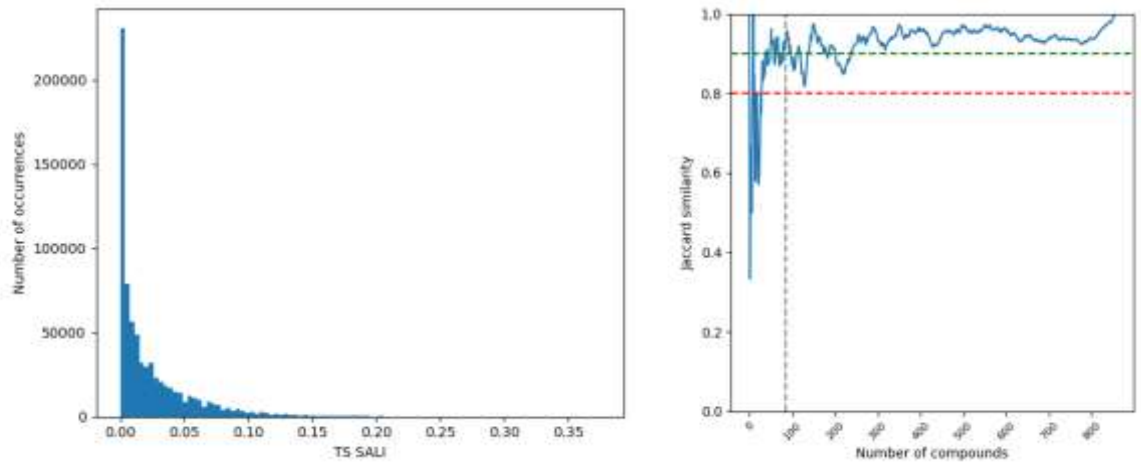


**Figure 21.** Distribution of pairwise TS_SALI values using 3rd order truncation. (Left) Variation of the Jaccard similarity between the ranking of cTS_SALI and complementary iCliff for database CHEMBL262_Ki. (right).

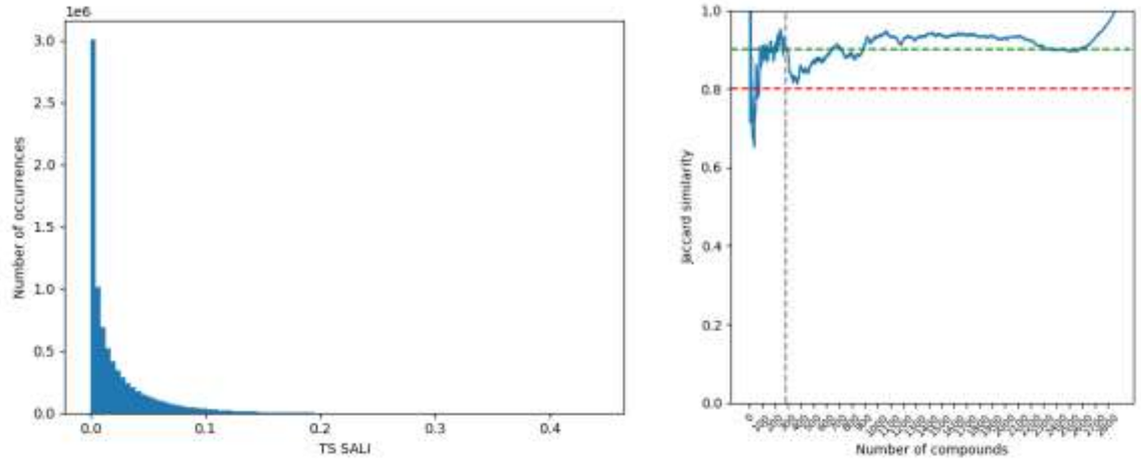


**Figure 22.** Distribution of pairwise TS_SALI values using 3rd order truncation. (Left) Variation of the Jaccard similarity between the ranking of cTS_SALI and complementary iCliff for database CHEMBL264_Ki. (right).

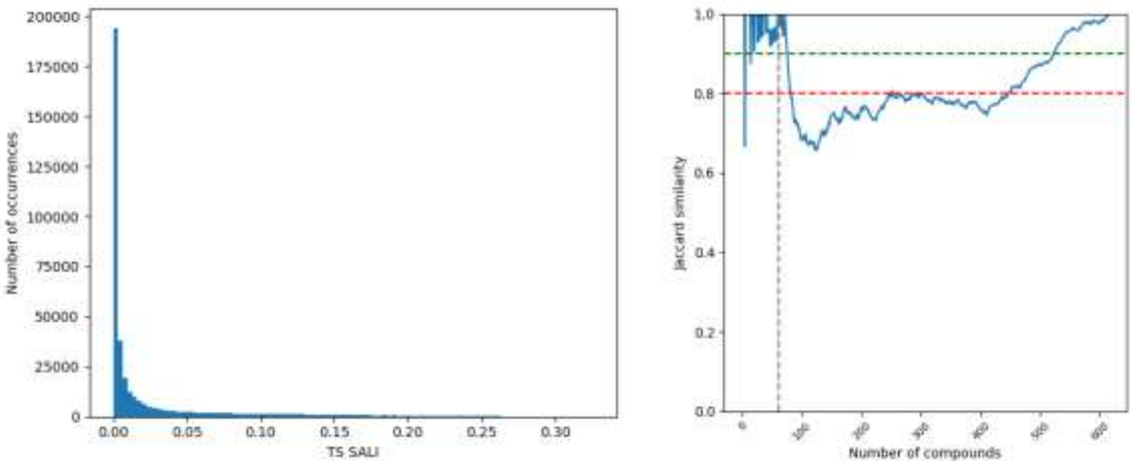


**Figure 23.** Distribution of pairwise TS_SALI values using 3$^{rd}$ order truncation. (Left) Variation of the Jaccard similarity between the ranking of cTS_SALI and complementary iCliff for database CHEMBL2835_Ki. (right).

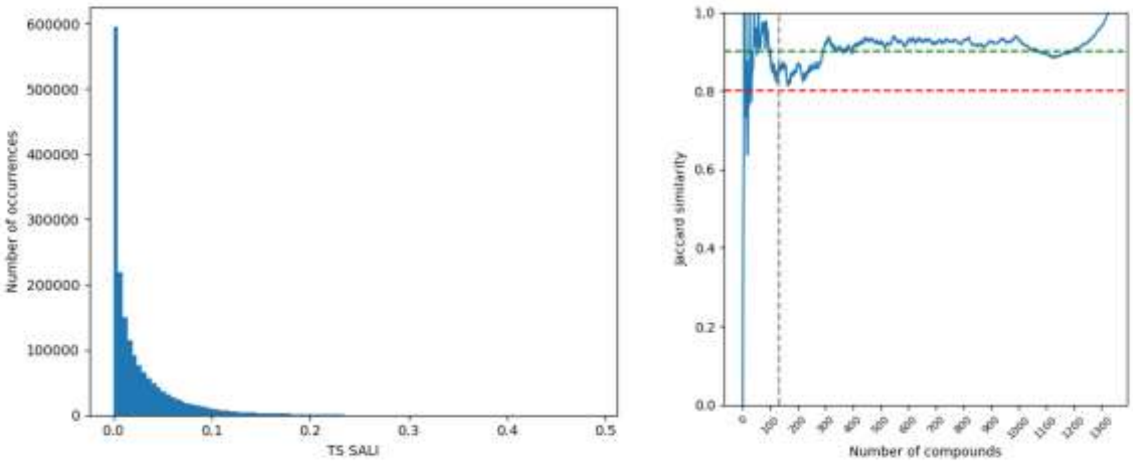


**Figure 24.** Distribution of pairwise TS_SALI values using 3$^{rd}$ order truncation. (Left) Variation of the Jaccard similarity between the ranking of cTS_SALI and complementary iCliff for database CHEMBL287_Ki. (right).

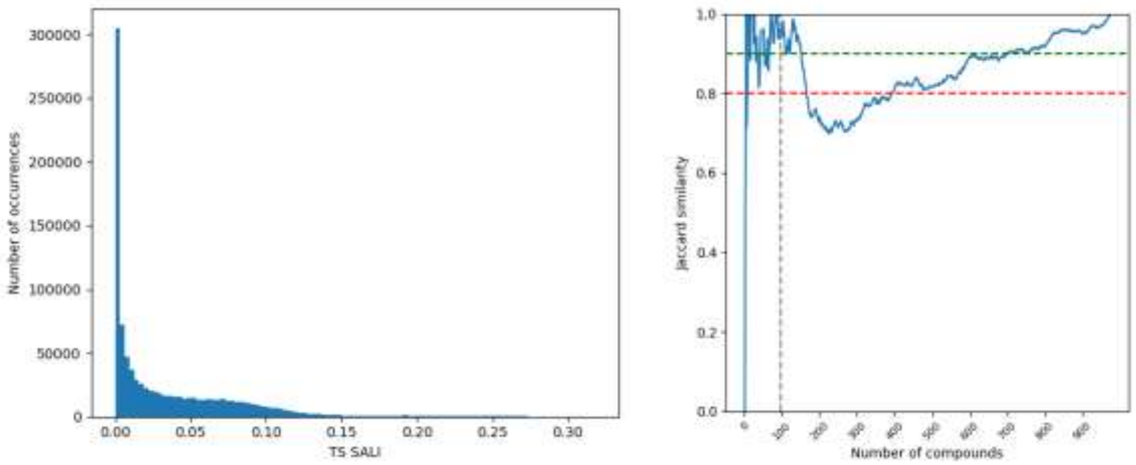


**Figure 25.** Distribution of pairwise TS_SALI values using 3rd order truncation. (Left) Variation of the Jaccard similarity between the ranking of cTS_SALI and complementary iCliff for database CHEMBL2971_Ki. (right).

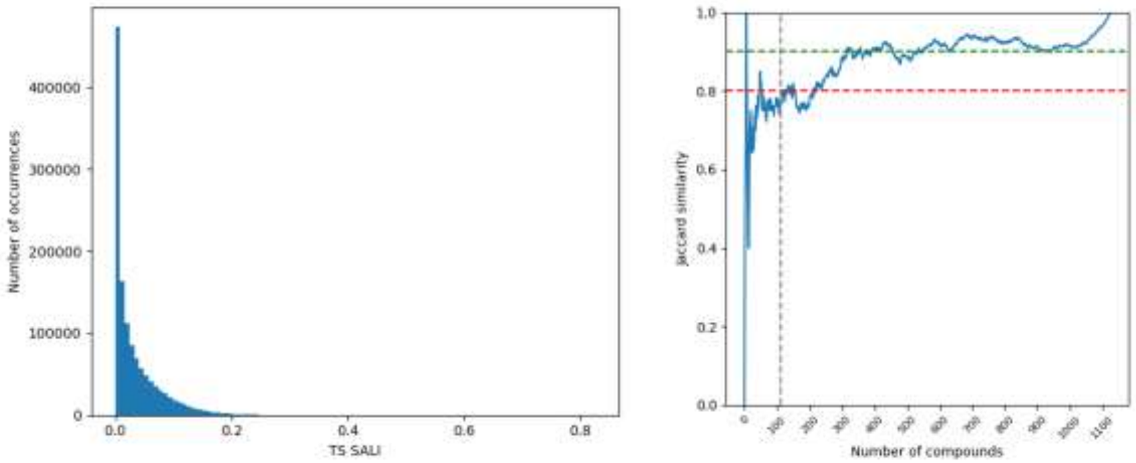


**Figure 26.** Distribution of pairwise TS_SALI values using 3rd order truncation. (Left) Variation of the Jaccard similarity between the ranking of cTS_SALI and complementary iCliff for database CHEMBL3979_EC50. (right).

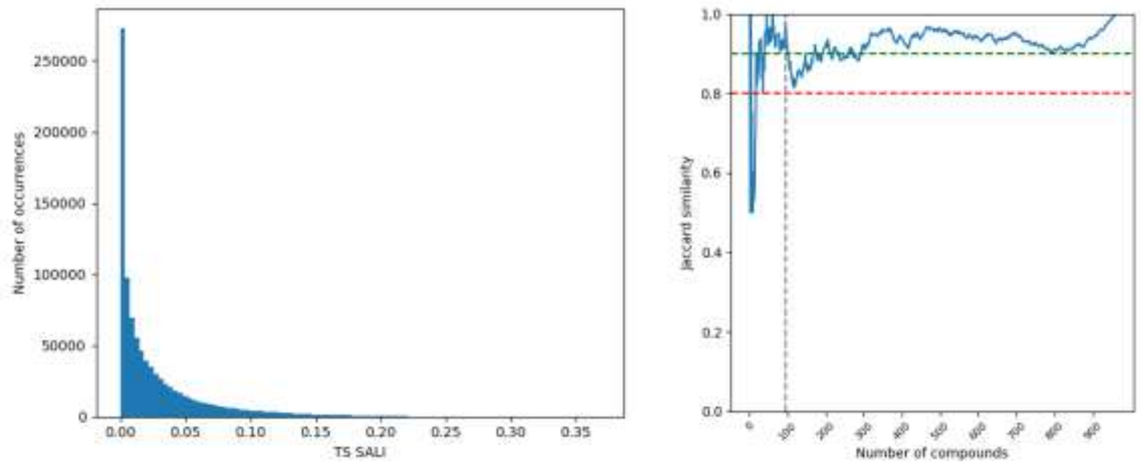


**Figure 27.** Distribution of pairwise TS_SALI values using 3rd order truncation. (Left) Variation of the Jaccard similarity between the ranking of cTS_SALI and complementary iCliff for database CHEMBL4005_Ki. (right).

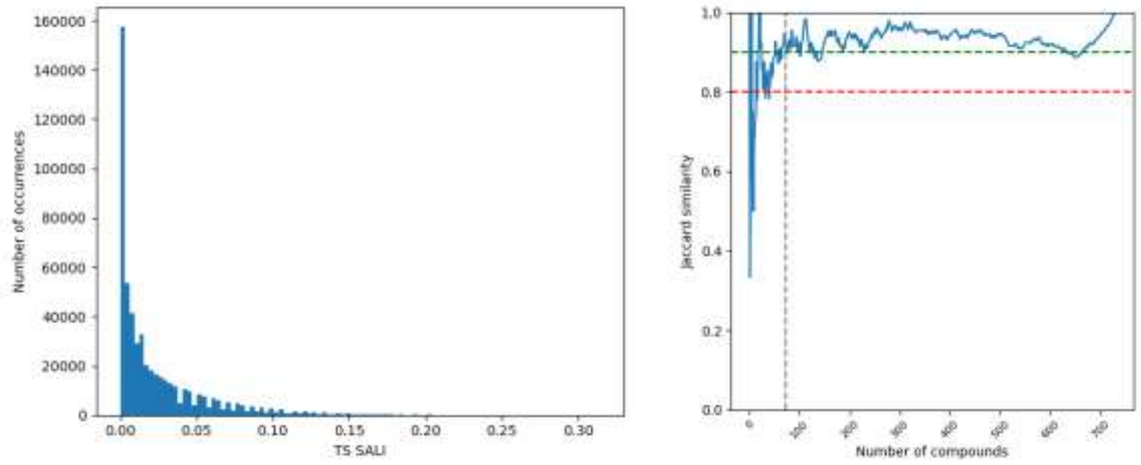


**Figure 28.** Distribution of pairwise TS_SALI values using 3rd order truncation. (Left) Variation of the Jaccard similarity between the ranking of cTS_SALI and complementary iCliff for database CHEMBL4203_Ki. (right).

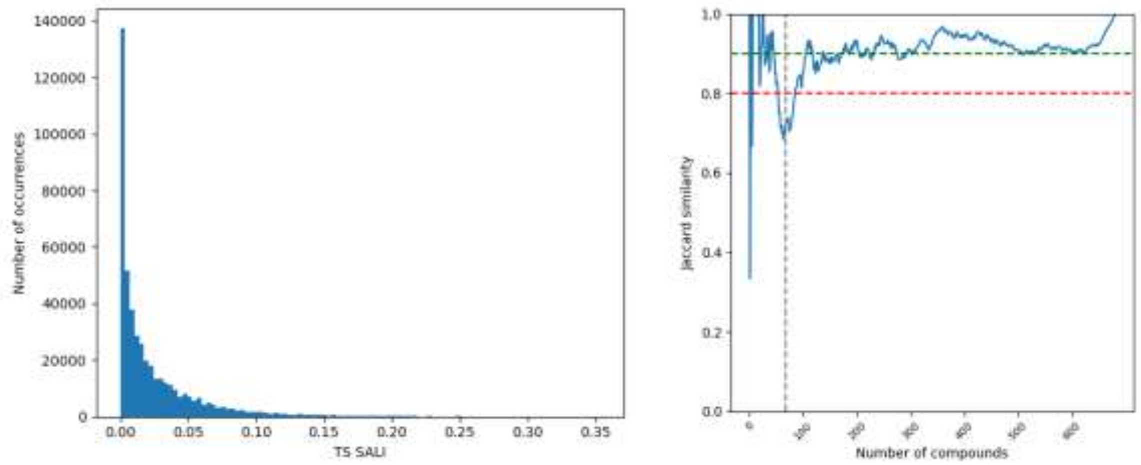


**Figure 29.** Distribution of pairwise TS_SALI values using 3rd order truncation. (Left) Variation of the Jaccard similarity between the ranking of cTS_SALI and complementary iCliff for database CHEMBL4616_EC50. (right).

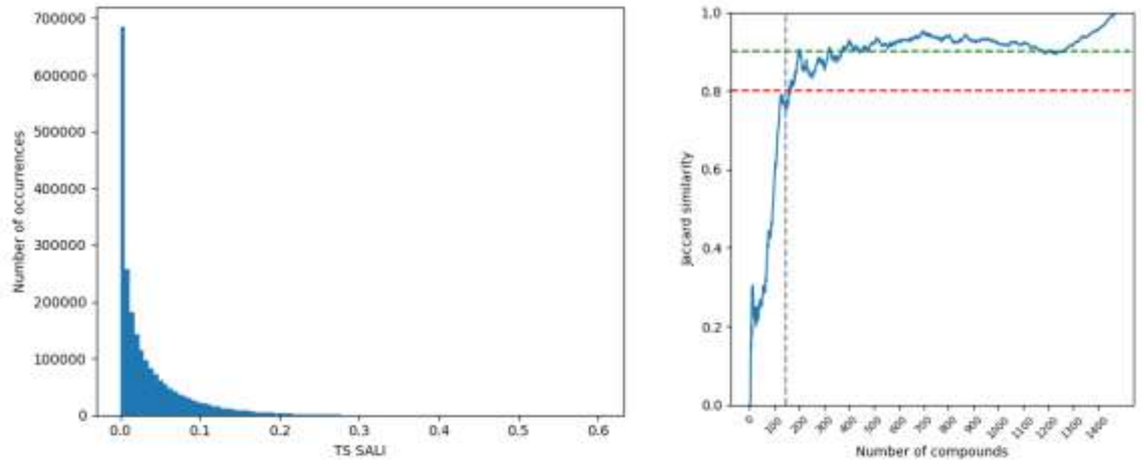


**Figure 30.** Distribution of pairwise TS_SALI values using 3rd order truncation. (Left) Variation of the Jaccard similarity between the ranking of cTS_SALI and complementary iCliff for database CHEMBL4792_Ki. (right).
